# Global reprogramming of virulence and antibiotic resistance in *Pseudomonas aeruginosa* by a single nucleotide polymorphism in the elongation factor-encoding gene, *fusA1*

**DOI:** 10.1101/842781

**Authors:** Eve A. Maunders, Rory C. Triniman, Taufiq Rahman, Martin Welch

## Abstract

*Pseudomonas aeruginosa* is a common opportunistic pathogen. The organism displays elevated intrinsic antibiotic resistance and can cause life-threatening infections. The gene encoding an elongation factor, FusA1, is frequently mutated in clinical isolates of *P. aeruginosa* from patients with cystic fibrosis (CF). Recent work has shown that *fusA1* mutants often display elevated aminoglycoside resistance due to increased expression of the aminoglycoside efflux pump, MexXY. In the current work, we isolated a spontaneous gentamicin-resistant *fusA1* mutant (FusA1^P443L^) in which *mexXY* expression was increased. Through a combination of proteomic and transcriptomic analyses, we found that the *fusA1* mutant also exhibited large-scale but discrete changes in the expression of key pathogenicity-associated genes. Most notably, the *fusA1* mutant displayed greatly increased expression of the Type III Secretion system (T3SS), widely considered to be the most potent virulence factor in the *P. aeruginosa* arsenal, and also elevated expression of the Type VI Secretion (T6S) machinery. This was unexpected because expression of the T3SS is usually reciprocally coordinated with T6S system expression. The *fusA1* mutant also displayed elevated exopolysaccharide production, dysregulated siderophore production, elevated ribosomal protein synthesis, and transcriptomic signatures indicative of translational stress. Each of these phenotypes (and almost all of the transcriptomic and proteomic changes associated with the *fusA1* mutation) were restored to levels comparable to that in the PAO1-derived progenitor strain by expression of the wild-type *fusA1* gene *in trans*, indicating that the mutant gene is recessive. Our data show that in addition to elevating antibiotic resistance through *mexXY* expression (although we also identify additional contributory resistance mechanisms), mutations in *fusA1* can lead to highly-selective dysregulation of virulence gene expression.

## Introduction

Due to its high intrinsic resistance to antibiotics and aggressive virulence, *Pseudomonas aeruginosa* holds the dubious accolade of consistently occupying a “top ten” slot on lists of clinical threats across the globe. Indeed, the World Health Organisation recently classified it as a top priority pathogen for which the development of new antimicrobial interventions is critical. This opportunistic, Gram-negative bacterium is ubiquitous and exhibits a particular predilection for the built environment, making encounters with the human populace commonplace. *P. aeruginosa* is frequently isolated from burn wounds, the respiratory tract, and the urinary tract, and is the leading cause of morbidity and mortality in people with cystic fibrosis (CF) (McCarthy *et al.*, 2014; Pereira *et al.*, 2014; Pang *et al.*, 2019). In CF, defective mucociliary clearance causes an accumulation of a thick mucus plugs within the airways. Such oxygen-limited environments provide the perfect niche for *P. aeruginosa* to thrive (Ratjen *et al.*, 2003; Trinh *et al.*, 2015).

Chronic *P. aeruginosa* infection of the CF lung is associated with the transition from an active, motile lifestyle to a sessile, biofilm-like mode of growth. These are bacterial communities embedded within a self-produced extracellular polymeric matrix, composed of mannose-rich polysaccharides, extracellular DNA and proteins (Maunders *et al.*, 2017). This matrix confers a level of protection against antibiotics and the host immune system (Alhede *et al.*, 2014). Biofilm formation is also associated with increased expression of the Type VI Secretion (T6S) machinery. The function of the *P. aeruginosa* T6S has become clearer in the last decade; it appears to play a role in killing other bacterial species (or even “non-self” *P. aeruginosa* strains) (Hood *et al.*, 2010; Basler *et al.*, 2013), especially in tightly packed biofilms where competition for the same resources is rife. Conversely, ‘free-swimming’ planktonic cells predominate in acute infection scenarios. Here, virulence factors and motility are up-regulated, and pathogenicity is enhanced (Breidenstein *et al*. 2011). Perhaps the most potent *P. aeruginosa* virulence determinant is the Type III Secretion System (T3SS), which mediates the translocation of cytotoxic effector proteins directly into the cytoplasm of neighbouring host cells.

These effectors subvert the function of the recipient cells, typically by disrupting the cytoskeleton, promoting cell rounding and apoptosis, and therefore assist immune evasion through preventing phagocytosis by host innate immune cells (Galle *et al.*, 2012; Berube *et al.*, 2017). Both the T3 and T6 secretion systems are contact-triggered injectosomes, however, they are structurally and mechanistically distinct, as are their targets, and the expression of these two systems appears to be inversely correlated (Moscoso *et al.*, 2011).

In previous work, we showed that the rapidly growing planktonic cells associated with increased virulence factor production display markedly up-regulated expression of the machinery required for macromolecular synthesis, especially proteins involved in translation (Mikkelsen *et al.*, 2007). Translation comprises four main steps; initiation, elongation, termination and recycling. The ribosome-associated protein, elongation factor G (EF-G, encoded by *fusA1* and *fusA2* in *P. aeruginosa*), is essential for two of these steps; ‘elongation’ and ‘recycling’ (Savelsbergh *et al*. 2009). During elongation, EF-G catalyses the translocation of charged tRNA from the A-site to the P-site, and from the P-site to E-site of the large ribosomal subunit. This is coupled with movement of the ribosome along the mRNA being translated. EF-G is comprised of five domains; domains I and II mediate GTP binding and hydrolysis, and domains III, IV and V dock to the A-site of the ribosome. The tip of domain IV interacts with the mRNA and promotes tRNA translocation (Salsi *et al.* 2015). This involves large structural rearrangements as the EF-G domains swivel relative to one another (Belardinelli *et al.*, 2017; Macé *et al.*, 2018). The translocation process is repeated until a stop codon is encountered and release factors catalyse hydrolysis of the peptidyl-tRNA bond, thereby liberating the newly synthesised polypeptide. EF-G then coordinates with ribosome recycling factor (RRF, a structural mimic of tRNA) to promote disassembly of the ribosomal subunits (Salsi *et al*. 2015; Wilson, 2014). Aminoglycoside antibiotics can disrupt both the elongation and recycling steps, thereby leading to a decrease in the overall number of ribosomes available.

Whole genome sequence analyses have revealed that *fusA1* is a hotspot for accruing mutations in *P. aeruginosa* isolates from patients with CF (Bolard *et al*, 2018). These *fusA1* mutants often display increased resistance to aminoglycoside antibiotics. However, little more is known about the phenotypic consequences of such *fusA1* mutations on the wider physiology of the cell. In this study we show that a spontaneous single nucleotide polymorphism (SNP) in *P. aeruginosa fusA1* gives rise to discrete, but large magnitude changes in key pathophysiological processes. For example, the strain containing the mutated *fusA1* allele displayed selective up-regulation of genes encoding the T3SS apparatus, the T6SS apparatus, exopolysaccharide biosynthesis genes, and a multidrug efflux system. These findings suggest a hitherto unexpected subtlety in the chain of events linking transcription, translation and virulence/antibiotic sensitivity in this organism.

## Results

### A SNP in *fusA1* causes decreased expression of the biofilm-associated protein, CdrA

At the outset of this investigation, we sought to identify potential regulators of biofilm formation in *P. aeruginosa*. To do this, we made a stable chromosomal reporter construct in which the promoter of the *cdrAB* operon (P*cdrAB*) was fused to a promoter-less *lacZ* ORF. CdrA is a biofilm-associated extracellular matrix adhesion, and is known to be primarily expressed in conditions that favour biofilm formation. The construct was integrated at a neutral site in the PAO1 chromosome using the mini-CTX system (Calvo *et al.*, 2000). The resulting P*cdrAB*::*lacZ* reporter strain, hereafter referred to as EMC0, yielded vivid blue colonies when grown on M9 minimal medium agar plates containing X-Gal and glucose, but far paler colonies on medium containing X-Gal and glycerol. This suggested that P*cdrAB* is activated during colony growth on glucose. To identify genes that might impinge upon transcription from the *cdrAB* promoter, we mutagenized EMC0 by introducing the pTn*Mod*-OGm plasposon (Dennis *et al.*, 1998). The resulting mutants were selected on plates containing gentamicin (to select for likely Tn insertion mutants) and X-Gal + glucose (to establish whether any of the mutants were affected in transcription from P*cdrAB*). One of the gentamicin resistant mutants that we isolated yielded colonies that exhibited a paler pigmentation than EMC0 on X-Gal/glucose plates, indicating a reduction in transcription from P*cdrAB* (**Figure 1A**). Further analysis of the mutant (hereafter, denoted EMC1) in M9-glucose liquid cultures confirmed the diminished β-galactosidase production. They also indicated that EMC1 had a minor growth defect in this medium (**Figure 1A**). We therefore measured growth and β-galactosidase production in a rich medium, LB. To our surprise, EMC1 exhibited an even greater growth defect in this medium (**Figure 1B**).

**Figure 1:**
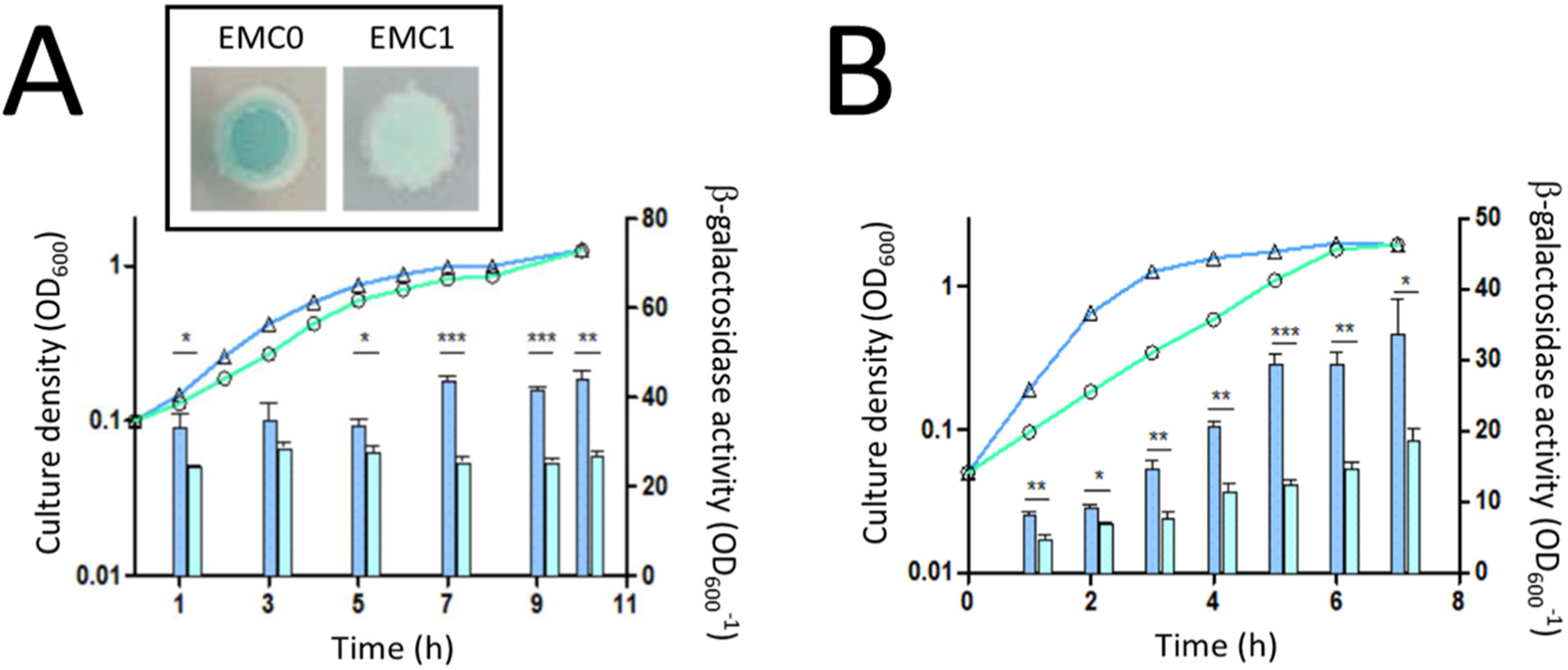
Transcriptional activity of the *cdrA* promoter was assessed by measuring β-galactosidase activity in the PAO1-derived reporter strain, EMC0, and in an EMC0-derived Gm^R^ mutant, EMC1. **(A)** β-galactosidase activity (bars) through the growth curves (lines) of EMC0 (blue bars and symbols) and EMC1 (light green bars and symbols) in M9 minimal medium containing 0.5% (w/v) glucose. *Inset*: Colonies of EMC0 and EMC1 grown on minimal media agar plates containing glucose and X-Gal. **(B)** β-galactosidase activity (bars) through the growth curves (lines) of EMC0 (blue bars and symbols) and EMC1 (light green bars and symbols) in LB. Statistical significance between the β-galactosidase activities of ECM0 and ECM1 was assessed by unpaired *t*-test (n=3) (* = *p* <0.05, ** = *p* <0.01, *** = *p* <0.001).

Attempts to identify the insertion site of the plasposon in EMC1 using conventional approaches (including “random primed” PCR-based amplification of the regions flanking the plasposon, or “cloning out” of the plasposon as previously described (Dennis *et al.*, 1998)) failed. We therefore used whole genome sequencing (WGS) of EMC1 (and, as a control, also of the EMC0 progenitor strain) to identify the plasposon insertion site. Remarkably, and in spite of the robust Gm^R^ phenotype of the strain (**Table 1**), we found that EMC1 did not contain a plasposon insertion. It did, however, contain a C**→**T transition at position 4,770,363 in the genome (**Figure S1**). This SNP was located in the *fusA1* ORF, and resulted in a proline to leucine substitution at position 443 in the protein. The SNP was confirmed by PCR amplification of the gene followed by Sanger sequencing of the PCR product. WGS also revealed a selection of additional potential SNPs in EMC1, but these were all subsequently found by PCR/Sanger sequencing to be false positive SNP calls arising from the WGS technology. We conclude that the only difference between EMC0 and EMC1 is the SNP in *fusA1*.

**Table 1.** Minimum inhibitory concentrations (MIC) of antibiotics for EMC0 and EMC1. MICs were determined in M9 minimal medium containing glucose as a sole carbon source after 24 hr growth.

| Antibiotic | Mode of Action | EMC0 | EMC1 |
| --- | --- | --- | --- |
|  |  | MIC (μg/mL) | MIC (μg/mL) |
| Gentamicin | Binds 30S ribosome and disrupts reading of tRNA | 20 | 150 |
| Kanamycin | Binds 30S ribosome and disrupts reading of tRNA | 1500 | >2000 |
| Tetracycline | Binds 30S ribosome and inhibits binding of tRNA | 20 | 20 |
| Chloramphenicol | Binds 50S ribosome and inhibits peptide bond formation | 250 | 250 |
| Rifampicin | Inhibits RNA polymerase | 100 | 100 |
| Fusidic acid | Inhibits elongation factor G | 1750 | 1750 |

The gentamicin resistance of EMC1 was heritable but somewhat unstable; progressive sub-culturing of the mutant in M9-glucose minimal medium lacking gentamicin gave rise to gentamicin-sensitive (Gm^S^) derivatives. After each 24 hr round of sub-culturing, aliquots were removed, serially-diluted, and plated onto non-selective M9-glucose agar. A selection of 40 colonies from these plates were then re-tested for gentamicin resistance by plating onto M9-glucose + Gm. Following the first round of sub-culturing (24 hr growth), none of the 40 tested colonies were Gm^S^. However, after the second round of sub-culturing (a cumulative 48 hr of growth) 1/40 colonies tested were gentamicin-sensitive, rising to 5/40 colonies after the third 24 hr round of sub-culturing. PCR-amplification and sequencing of the *fusA1* gene in these Gm^S^ derivatives revealed that they all still contained the *fusA1*^*P443L*^ mutation. This suggests that additional, second-site mutations are responsible for the reversion phenotype. These data also show that whatever gives rise to the Gm^R^ phenotype in EMC1 can be over-ridden by secondary mutations elsewhere.

### The proline→leucine substitution at position 443 in FusA1 affects protein stability and conformation

*FusA1* encodes one of the two paralogous elongation factor G (EF-G) proteins in *P. aeruginosa*, and plays a pivotal role in protein synthesis and ribosomal recycling (Palmer *et al.*, 2013). The amino acid sequence of the two paralogues (denoted *fusA1* (PA4266) and *fusA2* (PA2071)) is highly conserved, with a shared identity of 84%. EF-G1B, encoded by *fusA2*, is thought to have greater involvement in elongation and polypeptide synthesis (Palmer *et al.*, 2013). By contrast, EF-G1A (encoded by *fusA1*) has a more dominant role in ribosomal recycling and association with ribosomal recycling factors (Palmer *et al.*, 2013). FusA1 is a conformationally-flexible multi-domain protein (Lin *et al.*, 2015), and proline 443 sits in close proximity to the crucial GTPase “switch” regions (**Figure 2A**). The switch regions play an important role in directing GDP-GTP exchange, raising the question of whether the P443L substitution might affect the conformation of the protein. To test this, we measured the intrinsic tryptophan (Trp) fluorescence profile of purified wild-type FusA1 and FusA1^P443L^. Protein Trp fluorescence is exquisitely sensitive to the microenvironment of each Trp residue, and as such, can be a sensitive reporter of protein conformation. These analyses indicated that purified FusA1^P443L^ had a lower quantum yield at the Trp emission λ_max_ (332nm) compared with the wild-type protein (**Figure 2B**). This indicates that one or more Trp residues in the mutant protein are likely to exhibit altered solvent accessibility compared with the wild-type protein, possibly due to conformational differences. One of the more widely used web-based algorithms, mCSM (Pires *et al.*, 2014), predicted that the P443→L substitution should destabilise FusA1 by 0.267 kcal/mol. Consistent with this, purified FusA1^P443L^ had a lower melting temperature than the wild-type protein (**Figure 2C**). Taken together, these data indicate that the P443→L substitution likely alters the conformation, stability, or dynamics of FusA1.

**Figure 2.**
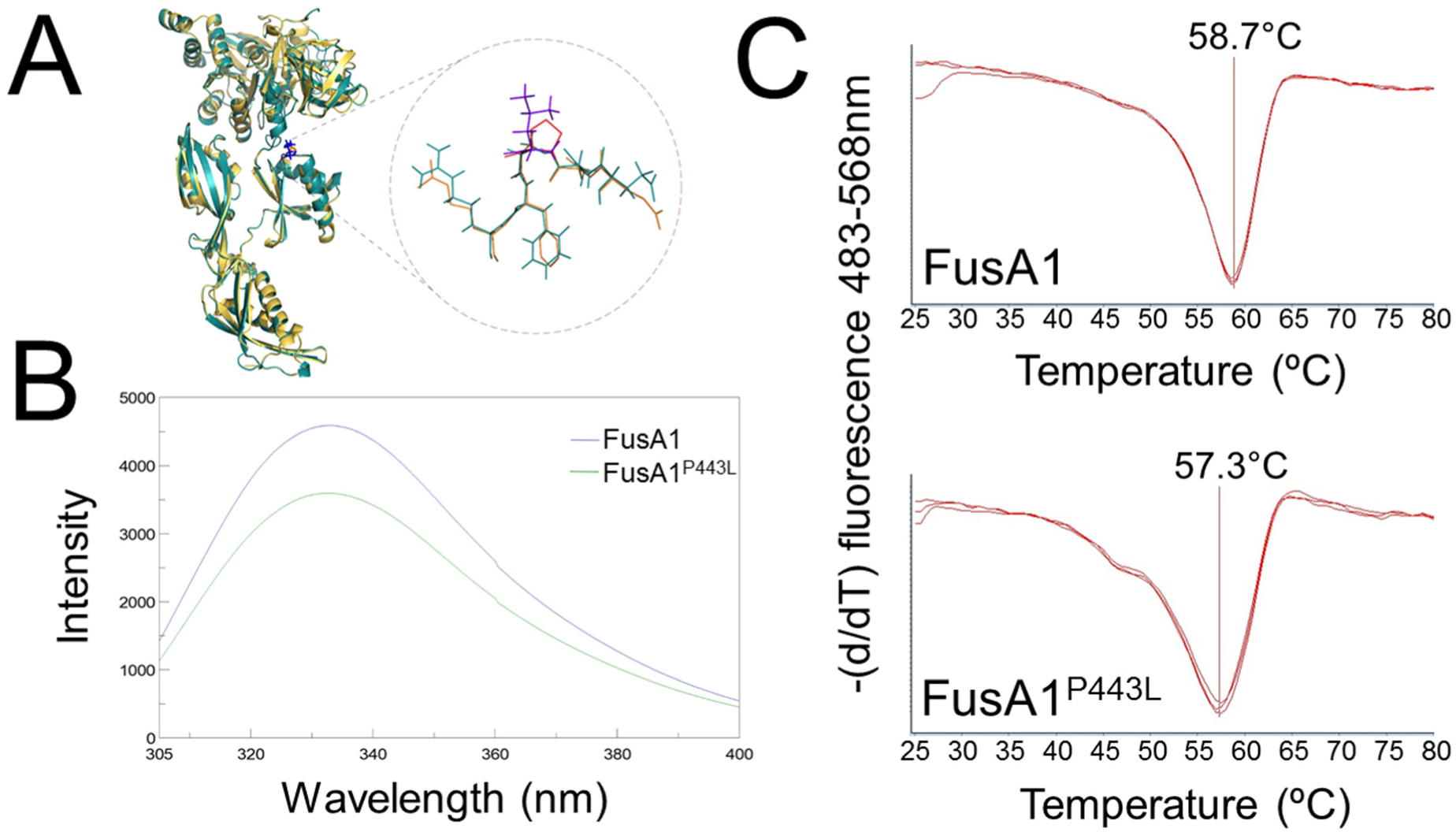
Structural impact of the P443L substitution on FusA1. **(A)** Location of the P443L substitution on the FusA1 structure. Proline 443 is highlighted in red. The “switch” regions important for GTP hydrolysis and domain organisation are indicated. **(B)** The tryptophan fluorescence emission spectrum of 0.8 μM purified wild-type FusA1 (blue line) and FusA1^P443L^ (green line) suggests that one or more Trp residues resides in an altered microenvironment in FusA1^P443L^. **(C)** Thermal shift data indicate that the wild type protein (upper panel) has a higher melting temperature than the FusA1^P443L^ protein (lower panel). This suggests that the P443L substitution decreases the thermal stability of the protein. Each melting curve was measured in triplicate.

### The P→L substitution in FusA1 has phenotypic consequences

The substantial growth defect associated with EMC1 grown in LB was complementable by expression of the wild-type *fusA1* gene *in trans* on a plasmid (**Figure S2**), suggesting that the wild-type allele is dominant. Compared with EMC0, EMC1 also exhibited defects in twitching motility and swimming motility, and these too could be complemented by introduction of wild-type *fusA1 in trans* (**Figure 3A**). EMC1 also displayed lower secreted gelatinase activity, and poor growth on the gelatinase plates. However, the lower secreted gelatinase production and slower growth of EMC1 may be linked, since complementation of EMC1 *in trans* with wild-type *fusA1* restored both phenotypes. Expression of *fusA1*^*P443L*^ *in trans* in EMC0 or EMC1 had no apparent effect on the motility/gelatinase phenotypes of the strains compared with the empty vector control. This suggests that the wild-type *fusA1* allele in EMC0 is dominant.

**Figure 3.**
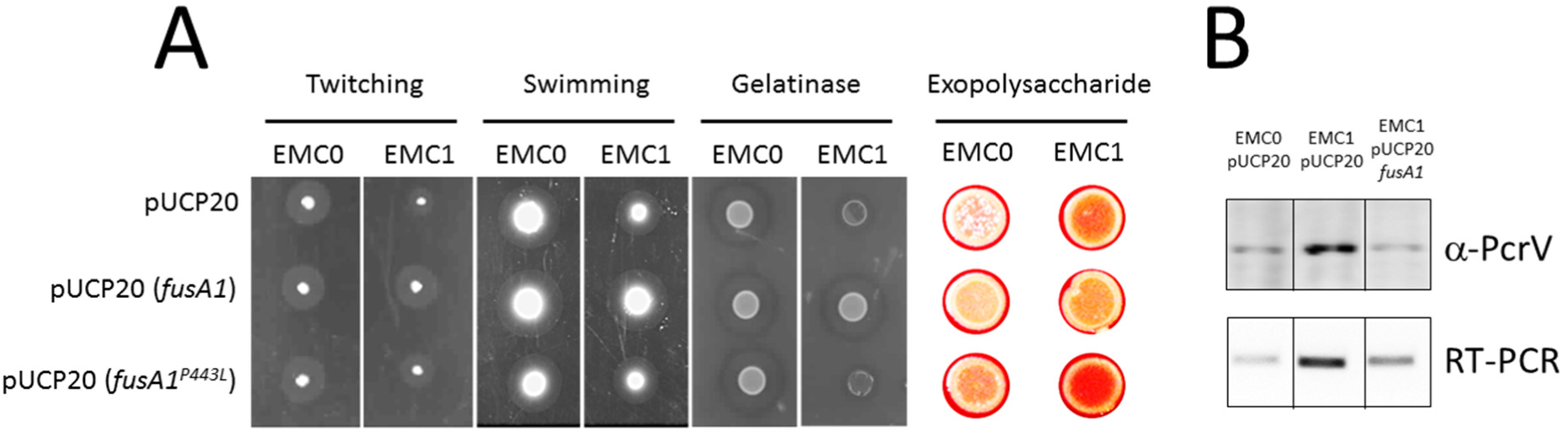
The *fusA1*^P443L^ mutation has pleiotropic effects on the cell. **(A)** The figure shows the twitching, swimming, gelatinase and exopolysaccharide phenotypes of EMC0 and EMC1 (as indicated) containing pUCP20 (empty vector control), pUCP20 encoding wild-type *fusA1*, or pUCP20 encoding *fusA1*^*P443L*^. Increased exopolysaccharide production is indicated by more intense staining of the colony with Congo Red. Expression of the *fusA1* mutant and wild-type ORFs cloned into pUCP20 was driven from the plasmid-encoded *lac* promoter. **(B)** Expression of PcrV protein (upper panel) and *pcrV*-encoding mRNA (lower panel) is increased in EMC1, and complemented by expression of wild-type *fusA1 in trans*. Representative results are shown. For replicates, see **Figure S4**.

By contrast with the impaired motility and secreted protease phenotypes, exopolysaccharide production was increased in EMC1 compared with EMC0, both on plate assays (**Figure 3A**) and in liquid culture (**Figure S3**). Expression of wild-type *fusA1 in trans* in EMC1 led to a decrease in exopolysaccharide production (compared with EMC1 containing the empty vector control), whereas expression of *fusA1*^*P443L*^ enhanced exopolysaccharide synthesis. Exopolysaccharides comprise the extracellular matrix which “glues together” cells in a biofilm. Interestingly in this regard, the increased exopolysaccharide production in EMC1 was not accompanied by an increase in its biofilm-forming ability compared with EMC0 (*data not shown*). Exopolysaccharide production is often inversely correlated with expression of the Type III Secretion (T3S) machinery. We therefore examined whether the P443L substitution in FusA1 impacted upon T3S. To our surprise, cultures of EMC1 over-expressed the Type III Secretion System (T3SS) protein, PcrV (**Figure 3B**). This increased expression was due to increased transcription of the *pcrV*-encoding operon, because RT-PCR analyses indicated that the amount of mRNA encoding *pcrV* was also increased in EMC1 (**Figure 3B** and **Figure S4**). This increased expression of PcrV could be reversed by supplying wild-type *fusA1 in trans*.

### Global consequences of the *fusA1*^*P443L*^ mutation on the proteome

Given that multiple phenotypes were affected by the *fusA1*^*P443L*^ mutation in EMC1, and that these phenotypes were not all modulated in the manner expected from previous studies (e.g., exopolysaccharide production and T3S both being up-regulated instead of inversely-regulated), this suggested that the P443L mutation may lead to global dysregulation in EMC1. To investigate this further, cultures of EMC0, EMC1 and EMC1 complemented with wild-type *fusA1* expressed from pUCP20 *in trans* (hereafter EMC1*) were grown to late exponential phase in M9 minimal media + glucose and were prepared for iTRAQ-based proteomic analysis. To establish whether any of the observed changes in protein profile were also underpinned by transcriptional changes, samples were also harvested for RNA-sequencing (from cultures grown in the same conditions).

The proteomic analysis resolved 3506 proteins (out of a total of 5570 predicted ORFs encoded by *P. aeruginosa* PAO1). Principal components analysis (PCA) of the data revealed that the proteome of EMC1 was distinct from that of EMC0, and that the proteomic changes giving rise to this segregation could be largely reversed by expression of wild-type *fusA1 in trans* in EMC1* (**Figure S5 A, S5B, S5C and Table S1**). Proteins were considered to be significantly modulated if they exhibited a log_2_-fold change (FC) > 1 (or < −1, if down-regulated) with a false discovery rate (FDR) adjusted P-value of ≤ 0.01. Based on these criteria, 128 proteins were up-regulated in EMC1 compared with EMC0, and 166 proteins were down-regulated. The 20 most highly up-regulated proteins are shown in **Table 2A**. Remarkably, and consistent with the earlier phenotypic analyses, over half (12/20) of these proteins were involved in T3S. Similarly, and also consistent with our earlier observations, PelA, involved in the biosynthesis of exopolysaccharide, was also up-regulated. A list of the top 20 down-regulated proteins is shown in **Table 2B**. The situation here is more ambiguous, since the majority (13/20) of these proteins currently have no assigned function.

**Table 2.**
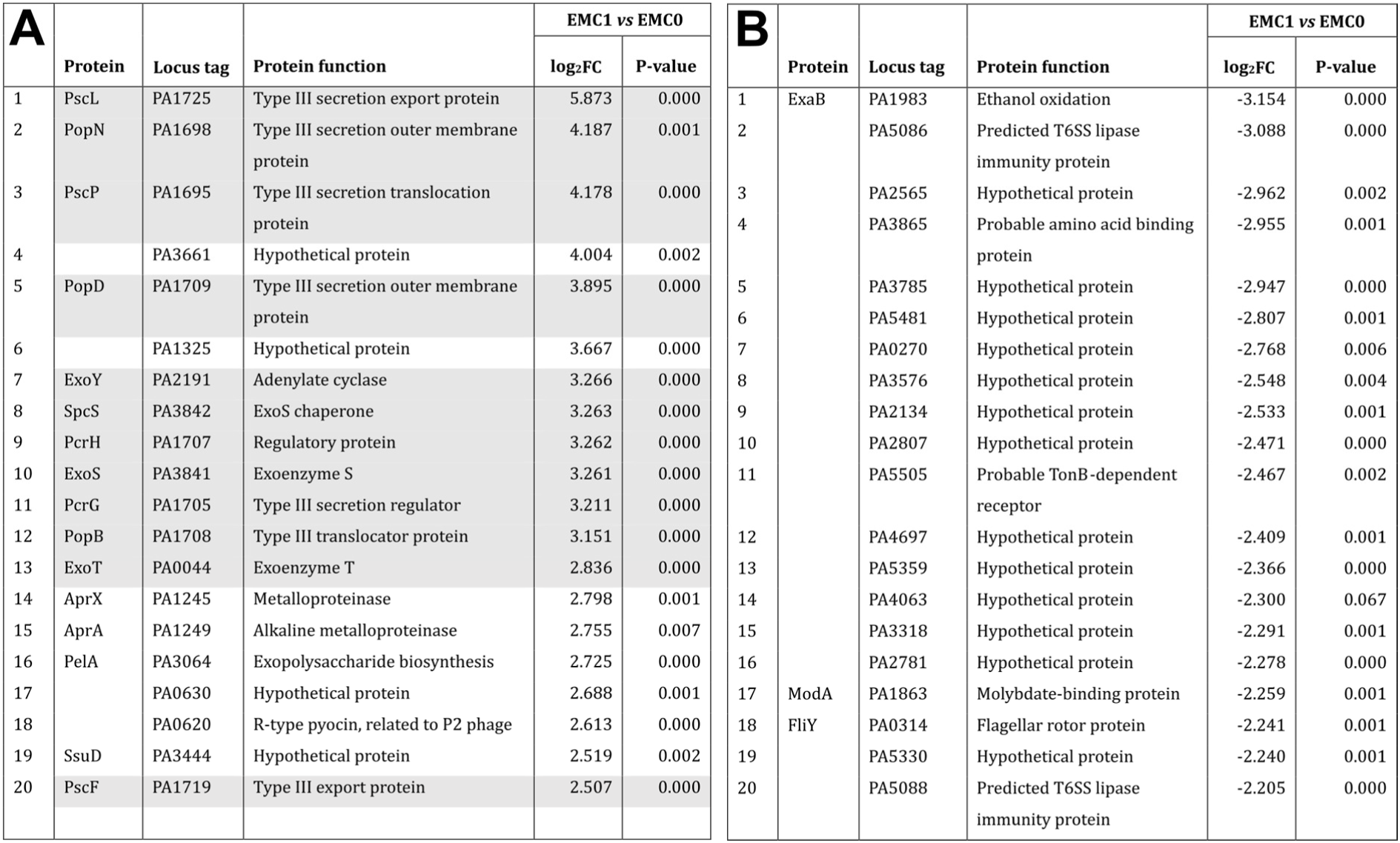
List of the top 20 proteins (based on log_2_FC) modulated in EMC1 *vs* EMC0. **(A)** Up-regulated proteins, rank-ordered by FC abundance. Proteins known to be involved in T3S are indicated by shading. **(B)** Down-regulated proteins, rank-ordered by FC abundance.

To obtain a more functional overview of the data, the 128 statistically-significantly up-regulated proteins in EMC1 were analysed using STRING to identify clusters of associated proteins. STRING is a database of known and predicted physical- and functional protein-protein interactions (Szklarczyk *et al.*, 2019). Inspection of the STRING output revealed that the modulated proteins fell into several distinct functional clusters involved in a variety of pathophysiological processes. The most obvious cluster comprised proteins associated with the T3SS (**Figure 4**). The likely driver behind this was the ca. 2-fold up-regulation of ExsA, which is the master regulator (activator) of T3SS expression (Diaz *et al.*, 2011). Also consistent with the phenotypic data, a cluster of proteins (PelA, PelF and PslG) involved in exopolysaccharide biosynthesis were up-regulated in EMC1. The data also revealed a probable explanation for the enhanced gentamicin resistance of EMC1; ArmZ, a major activator of the *mexXY* aminoglycoside efflux pump, was up-regulated, as was expression of MexXY (Morita *et al.*, 2012; Hay *et al.*, 2013). Unexpectedly, we also noted that a selection of proteins involved in biosynthesis of the siderophore, pyochelin, were significantly up-regulated. This was somewhat surprising since (i) proteins associated with the biosynthesis of the other major siderophore in *P. aeruginosa*, pyoverdine, were unaffected, (ii) the abundance of iron-uptake master regulators, Fur and PvdS, as well as the pyochelin-specific regulator PchR, were unchanged in EMC1 compared with EMC0, and (iii) phenotypic analyses revealed that cultures of EMC1 produce less secreted siderophore(s) than EMC0 or EMC1* (**Figure S6**). It therefore seems that in EMC1, the biosynthetic pathway for pyochelin is expressed, but the siderophore is not secreted. This dysregulation of iron homeostasis in EMC1 may have additional consequences. A link between iron availability and T3S has been documented in several bacterial genera including *Bordetella, Salmonella, Shigella, Edwardsiella, Vibrio* and *Yersinia* (Wilderman *et al.*, 2004; Bronstein *et al.*, 2008; Gode-Potratz *et al.*, 2010; Chakraborty *et al.*, 2011; Kurushima *et al.*, 2012; Miller *et al.*, 2014). To investigate further whether iron availability impacts upon expression of the T3SS in EMC1, we examined whether supplementation of the growth media with additional iron had any effect on PcrV expression. The additional iron had little effect on expression of PcrV in EMC0 or EMC1*. However, addition of excess iron to the EMC1 cultures led to increased PcrV expression (**Figure S6**). It is therefore possible that the dysregulation of iron homeostasis in EMC1 is another factor that impacts on expression of the T3SS.

**Figure 4.**
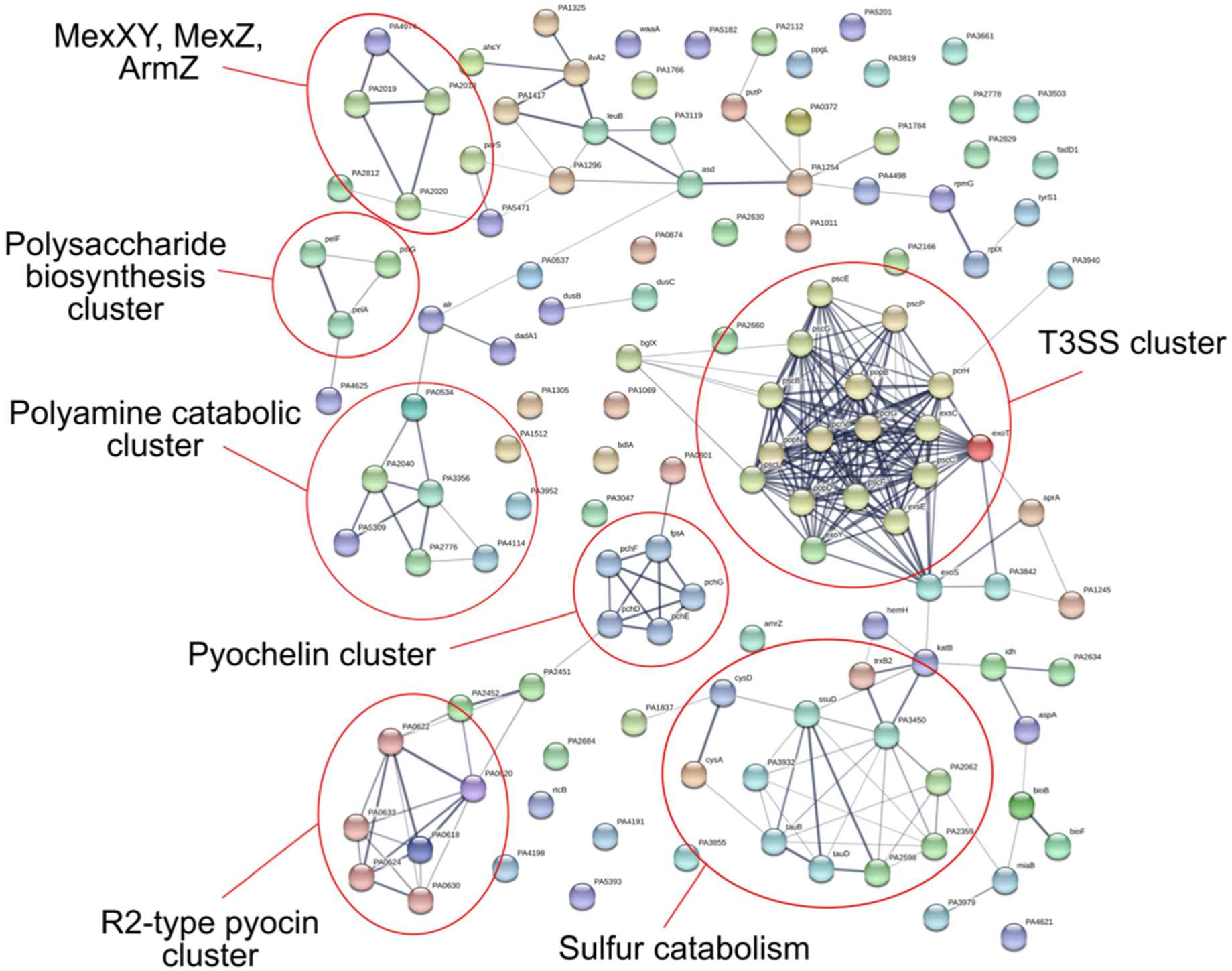
STRING-based analysis of up-regulated proteins in EMC1. The indicated clusters are discussed in the main body text.

A number of bacteriophage-like R2-type pyocin gene products were also up-regulated. R-type pyocins cause membrane depolarisation and inhibit active transport in closely related species to reduce bacterial competition (Llamas *et al.*, 2007). Another obvious functionally-related grouping was comprised of proteins associated with sulfur metabolism (including CysA, CysD, SsuD, TauA and TauB). However, perhaps the most notable cluster of up-regulated metabolic proteins were those involved in polyamine catabolism (PauA4, PauA5, PauB1, PauB3, PauB4). The genes encoding these proteins are strongly induced by the polyamines, putrescine and spermidine, and are distributed across the chromosome (Chou *et al.*, 2013). These gene products encode probable γ-glutamylpolyamine synthases involved in converting putrescine into γ-aminobutyric acid, and their collective up-regulation indicates that putrescine and/or spermidine may be more abundant in cultures of EMC1. Consistent with this notion, spermidine is known to also increase the expression of the T3SS proteins (Zhou *et al.*, 2007)

We also carried out a STRING-based analysis of the proteins that were down-regulated in EMC1. Fewer clusters were apparent compared with the up-regulated STRING map. This likely reflects the large number of uncharacterised proteins in the dataset; with little information to functionally link these proteins, clusters will inevitably be more sparsely populated. However, some patterns were apparent. From the STRING map (**Figure S7**), a small cluster comprising ExaB, nitrite reductase (NirS), and azurin (Azu) is discernable. ExaB is of interest because it was the protein with the greatest FC (**Table 2B**). ExaB is a cytochrome C550 involved in the breakdown of ethanol to aldehyde (Schobert *et al.*, 1999). Azurin transfers electrons from another c-type cytochrome, CytC551, to cytochrome oxidase. Cytochrome C551 is also the electron donor of nitrite reductase (NirS) and plays a role in dissimilative denitrification (Cutruzzolà *et al.*, 2002). These findings suggest that elements of the bacterial electron transport chain may be affected in EMC1. The ExaB/NirS/Azu cluster was linked to an adjacent group of proteins with roles in disulphide bond formation (DsbA, DsbD2, a probable glutathione peroxidase (PA1287), TrxA, and three other Trx-like proteins (PA0941, PA3664, PA0950)) and protein folding (PpiA, PpiC2). These proteins are closely linked to the electron transfer chain and cellular redox status.

Several other highly down-regulated proteins were linked to the T6SS. PA5086 and PA5088 are predicted to encode T6S lipase immunity proteins, protecting the producing cell from lipase inflicted self-harm (Dong *et al.*, 2013). PA5088 is operonic with an encoded T6S phospholipase D effector (*tleB5*) and with *vgrG5*, although neither of these proteins were affected in our experiments. VgrG proteins are secreted by the T6SS to form complexes that perforate host cell membranes, and the genomic position of *vgrG5* adjacent to PA5088 makes it likely that its expression was also reduced. The down-regulation of T6SS-assocated proteins observed here is consistent with the known inverse regulation of T6S and T3S (Moscoso *et al.*, 2011). Another discernible cluster of proteins was related to cell motility and chemotaxis, including CheR1, PA0177, PctC, PA1464, and the twitching motility protein, PilH. The flagellar motor protein, FliY, and an uncharacterised protein, PA2781, were also down-regulated. PA2781 is likely operonic with PA2780 (*bswR*), a bacterial swarming regulator (undetected in the proteomic analysis), and is predicted to interact with FlgB, a structural component of the bacterial flagellum. This suggests that PA2781 may also affect flagella-mediated motility. Collectively, these observations are consistent with the decreased twitching and swimming motility associated with EMC1 (**Figure 3**).

### Global consequences of the *fusA1*^*P443L*^ mutation on the transcriptome

Given the role of FusA1 in translation, and given the apparently very selective consequences of the *fusA1*^*P443L*^ mutation on the proteome, we wondered whether these changes might also be reflected at a transcriptomic level too. To address this further, we carried out an RNA-Seq analysis of the mRNA expression profiles in EMC1 and EMC1* compared with EMC0. Of the 5678 predicted RNA-encoding genes in *P. aeruginosa* PAO1, 5628 were represented in the RNA-Seq dataset. This suggests that most ORFs are expressed at detectable levels in the conditions tested. Of these expressed genes, 657 were significantly (P < 0.01) up-regulated (FC > 1), whereas 374 were significantly down-regulated (FC > −1) when comparing EMC0 with EMC1. Far fewer transcripts were significantly modulated when comparing EMC0 with EMC1* (**Figure S8**), indicating that expression of the wild-type *fusA1* gene was able to complement many of the changes in EMC1. This was also reflected by the lower level of scatter away from the midline in the FPKM (fragments per kilobase, per million) plots (**Figure S8**).

The top 20 most highly up-regulated transcripts are shown in **Table 3A**. Most (16/20) of these are involved in T3S. This indicates that the T3SS is up-regulated (by up to 46-fold) at a transcriptional level as well as at a translational level in EMC1. These changes were reversed back towards the levels seen in EMC0 by the introduction of the wild-type *fusA1* gene *in trans* (**Table S1**).

**Table 3.**
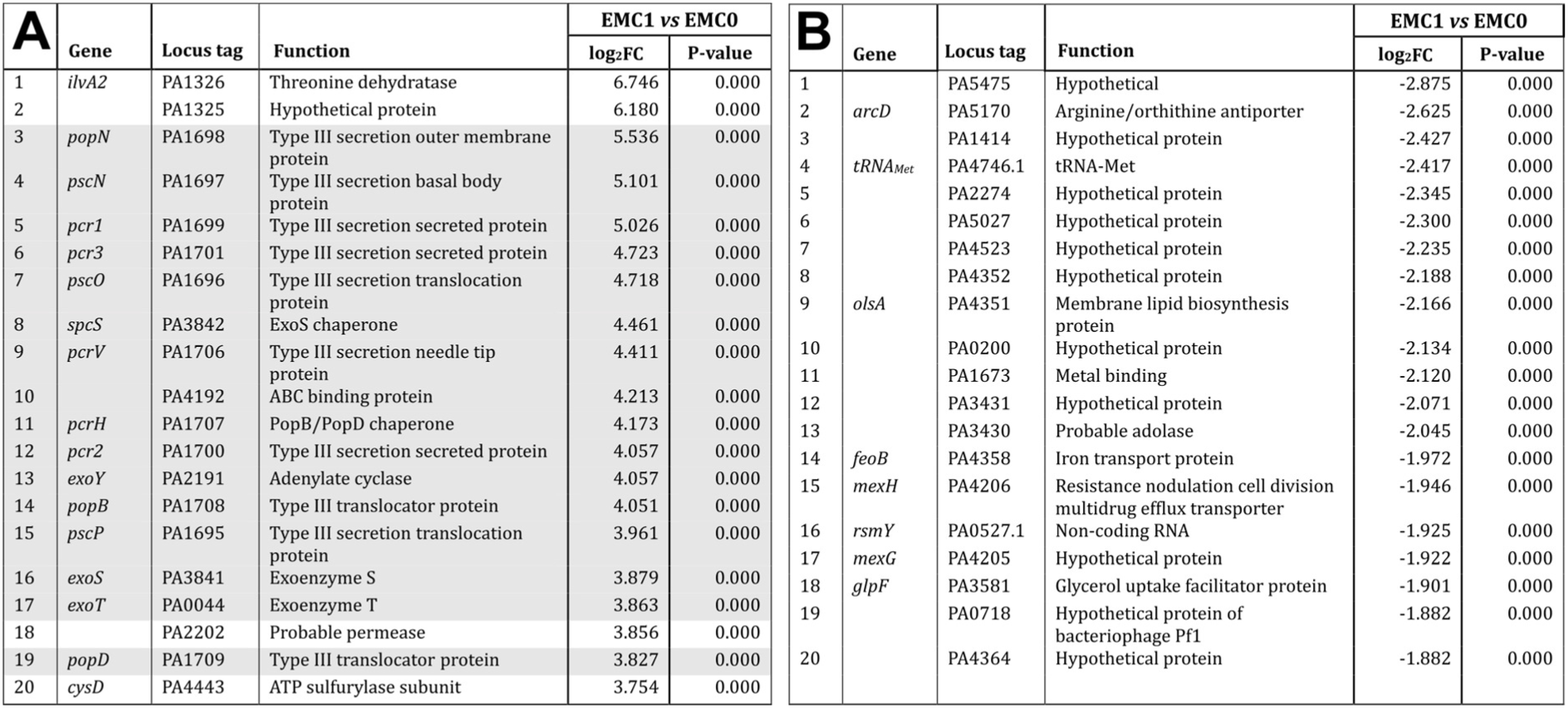
List of the most highly up-regulated transcripts (A) and down-regulated transcripts (B) associated with the *fusA1*^*P443L*^ mutation. Shaded boxes in **(A)** feature genes associated with the T3SS. A P-value of 0.000 represents < 0.0005.

To establish potential functional interactions between the modulated transcripts, we carried out a STRING interactome analysis. Due to the large number of significantly modulated transcripts, only the top 200 were used to predict interactions between the encoded proteins. T3SS genes were by far up the largest cluster on the STRING protein interaction map (**Figure 5**). Consistent with the proteomic analysis, *exsA* expression was up-regulated by around 3-fold, and is most likely to be a major factor driving the observed global up-regulation of the T3SS transcripts. It is worth noting that the transcriptional regulator PsrA is thought to be required for full activation of the *exsCEBA* operon (Shen *et al.*, 2006). However, *psrA* transcripts exhibited a 2-fold reduction in expression, suggesting that PsrA may not be a major requirement for T3S in all conditions, and that low levels of *psrA* transcription do not necessarily prevent expression of the *exsCEBA* operon. Other distinct clusters predicted by STRING included a group of ribosome-associated proteins, and surprisingly, also a robust cluster of T6SS-associated transcripts (mostly from Hcp Secretion Island I (HSI-I)). Indeed, the transcript encoding the HSI-I needle protein, Hcp1, was up-regulated 13-fold, and many other T6-associated transcripts were up-regulated >4-fold. This was unexpected because the T3SS and T6SS are reciprocally-regulated in most conditions, and several T6S-associated proteins were down-regulated in the proteomic analyses.

**Figure 5.**
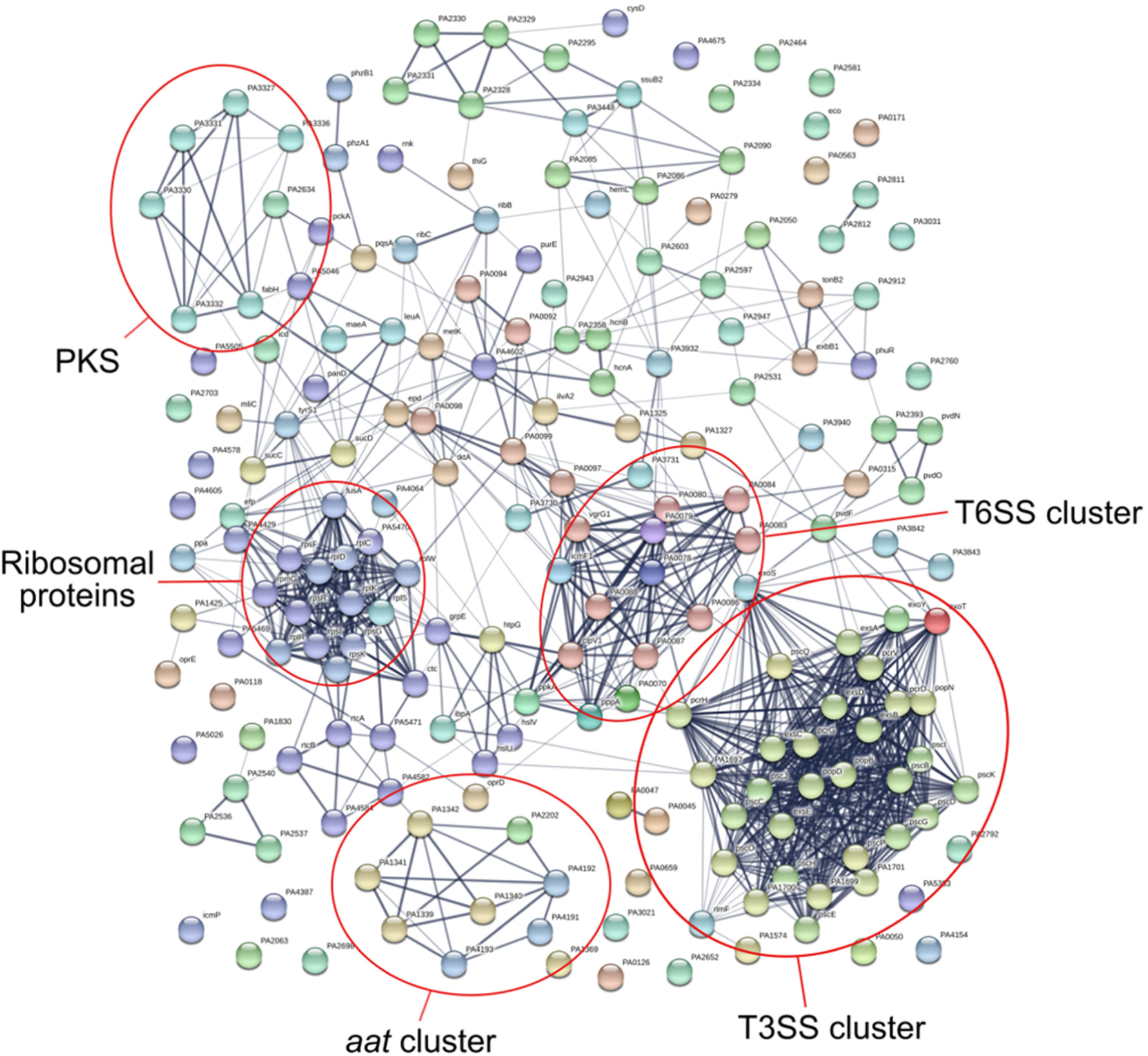
STRING-based analysis of up-regulated transcripts in EMC1. The indicated clusters are discussed in the text.

Although the list of most-highly up-regulated transcripts was dominated by the T3SS genes, the two genes with the highest modulation in EMC1 were an operonic pair, PA1325 and PA1326 (**Table 3**). PA1326 (*ilvA2*) encodes a threonine dehydratase involved in glycine, serine and threonine metabolism, and the adjacent gene, PA1325, is uncharacterised. Indeed, a relatively large number of genes linked to amino acid metabolism and were found to be up-regulated in this region of the genome. For example, the nearby *ggt* (PA1338) gene encodes a γ-glutamyltranspeptidase involved in glutathione catabolism (up-regulated in EMC1 2.6-fold). Adjacent to *ggt* is another set of up-regulated genes encoding an ABC transport system for glutamate and aspartate (*aatP* (PA1339, ↑4.0-fold), *aatM* (PA1340, ↑4.5-fold), *aatQ* (PA1341, ↑6.1-fold) and *aatJ* (PA1342, ↑3.7-fold), (Kahlon, 2016)). Another up-regulated gene was PA3965 (↑2.4-fold), encoding an AsnC-type transcriptional regulator which, although uncharacterised, is one of just two Lrp homologs encoded by PAO1. In *P. aeruginosa*, Lrp regulates the expression of genes involved in amino acid biosynthesis and catabolism (Diraviam Sriramulu, 2009). We also noted that transcripts from an uncharacterised gene cluster (PA3327-PA3336) encoding a non-ribosomal peptide synthase were up-regulated. The product of the non-ribosomal peptide synthase in the cluster (PA3327, ↑6.6-fold) was predicted by antiSMASH to generate a dipeptide core based around the condensation of serine and proline. Another large cluster (comprising 62 up-regulated transcripts in all) encoded ribosomal proteins and translation factors. Of these, 20 encoded 30S subunit proteins, and 26 encoded 50S subunit proteins. Peptide chain release factors, elongation factor Tu and initiation factor-2 were also up-regulated. One possible reason for the overall up-regulation of ribosomal transcripts in EMC1 came from the earlier proteomic analysis, which revealed increased abundance of SmpB. This is an RNA-binding protein that is induced upon ribosomal stalling. The increased expression of SmpB indicates that the mutated FusA1 likely leads to decreased translation efficiency, and is compensated for by an increase in the overall expression of the translation machinery. Several genes involved in RNA processing were also up-regulated, including the RNA 3’-terminal phosphate cyclase (*rtcA*, ↑5.2-fold) and RNA ligase (*rtcB*, ↑9.3-fold).

A large number of transcripts were also down-regulated in EMC1, although generally, the magnitude of modulation was low (**Table 3B**). A STRING analysis of the 200 most highly down-regulated transcripts is shown in **Figure S9**. Rather few distinct clusters were apparent; again, this is likely due to the predominance of transcripts from uncharacterised genes in the dataset. A few of the identified genes are worth commenting upon though. Firstly, the small untranslated RNA, rsmY (PA0527.1) was down-regulated 3.8-fold. The rsmY binding partner and antagonist, RsmA, was also down-regulated (↓5.7-fold). These are potentially important observations because excess “free” (i.e., unsequestered, by rsmY) RsmA increases expression of the T3SS genes and depresses expression of the T6SS/exopolysaccharide genes (Hood *et al.*, 2010). The expression of *rsmY* is thought to be regulated by the Gac signalling pathway. However, none of the other Gac/Rsm pathway-encoding transcripts (*ladS, retS, gacA, gacS*) were modulated in EMC1, nor was rsmZ (another untranslated small RNA thought to act in a similar manner to rsmY). If current models are correct, a stoichiometric imbalance of rsmY:RsmA levels should impinge reciprocally on T6SS expression and extracellular polysaccharide production (on the one hand), and T3SS expression (on the other). That we saw an *increase* in the expression of T3SS, T6SS and exopolysaccharide biosynthetic genes suggests that signalling through the Rsm pathway may be dysregulated in EMC1. Secondly, and consistent with the proteomic observations, *mexXY* expression was up-regulated at a transcriptional level (↑≥2-fold). EMC1 exhibited down-regulation of *pprA* expression (↓2.4-fold). PprA and PprB are predicted to be part of a two-component regulatory system controlling membrane permeability. *PprAB* expression leads to increased membrane permeability and increased sensitivity to antibiotics, including aminoglycosides (Wang *et al.*, 2003). The decreased expression of *pprA* may have been enough to prevent activation of the two-component system and to decrease cell permeability, thereby also contributing towards the elevated aminoglycoside resistance in EMC1. Another factor that impinges upon *mexXY-oprM* expression is the extracytoplasmic sigma factor, SigX (Gicquel *et al.*, 2013). *SigX* transcripts exhibited a slight decrease (↓1.3-fold) in expression in EMC1. This is potentially significant because SigX normally stimulates expression of the response regulator, PprB, and Gicquel *et al.* (2013) have also reported elevated *mexXY* expression in a *sigX* mutant. Taken together, the decreased expression of both *sigX* and *pprA*, as well as the increased expression of the *mexXY* transcriptional activator, ArmZ (observed in the proteomics) likely explains the elevated gentamicin resistance of EMC1 (**Table 1**). The transcriptomic data also revealed that, in contrast with the observed up-regulation of *mexXY*, another RND-family efflux pump, *mexGHI-opmD*, was down-regulated in EMC1. This efflux system provides resistance to a variety of xenobiotics, and has recently also been associated with binding a quorum sensing molecule, the Pseudomonas Quinolone Signal (PQS) (Hodgkinson *et al.*, 2016). Although the precise physiological function(s) of MexGHI-OpmD remain to be elucidated, it has no known association with gentamicin resistance (Sekiya *et al.*, 2003; Sakhtah *et al.*, 2016). Finally, and consistent with the original aim of the study, *cdrA* transcription was down-regulated (↓1.4-fold) in EMC1.

## Discussion

In this work, we serendipitously identified a SNP in the *fusA1* gene of *P. aeruginosa* which gave rise to increased gentamicin resistance and global changes in expression of the T3 and T6 (HSI-I) secretion systems and exopolysaccharide biosynthetic pathways. *FusA1* mutations are a common feature in certain clinical *P. aeruginosa* isolates (such as those derived from CF sputum), and appear to be a low-cost response to exposure to sub-lethal concentrations of aminoglycosides *in vitro* (Chung *et al*., 2012; Mogre *et al.*, 2014; Bolard *et al.*, 2018). However, to our knowledge, the wider phenotypic consequences of such mutations have not been explored further. The current work shows that at least one such *fusA1* mutation can have selective, but potentially clinically-significant effects beyond conferring antibiotic resistance, by affecting the expression of key virulence factors. Early indications that the *fusA1*^*P443L*^ mutant (EMC1) might be pleiotropically affected came from the unexpected observation that it had a larger growth defect (compared with the wild-type progenitor) in rich media than it did in minimal media. Our transcriptional data – showing that ribosomal gene expression is increased in EMC1 - indicate that this is likely caused by the increased (but unmet, due to defective FusA1^P443L^ function) translational demand during rapid growth in rich medium.

The elevated gentamicin resistance associated with EMC1 and other *fusA1* mutants (Bolard *et al.*, 2018) appears to be driven by increased expression of the *mexXY*-encoded aminoglycoside efflux pump. This, in turn, is likely due to increased expression of the cognate transcriptional activator of *mexXY* expression, ArmZ. In addition, EMC1 also displayed transcriptional hallmarks indicative of decreased cell envelope permeability to aminoglycosides (linked to lower *sigX* and *pprA/B* expression). It seems highly unlikely that the *fusA1* mutation itself contributes directly towards gentamicin resistance because “bypass” mutants in which gentamicin sensitivity is restored arise at high frequency. These appear to be due to the acquisition of second-site mutations because the *fusA1*^*P443L*^ mutation is retained in all of the cases tested. That there is a selection pressure to acquire such bypass mutations indicates that the changes which accompany gentamicin resistance have a fitness cost. Current efforts are aimed at identifying the gene(s) which acquire these bypass mutations.

Interestingly, post-translational modifications (such as those carried out by ErmBP or TetO) which prevent the binding of antibiotics to the ribosome, or resistance-conferring mutations which block the binding of antibiotics to the ribosome, also prevent induction of *mexXY* (Jeanott *et al*., 2005). Furthermore, although most of the antibiotics that are known to induce *mexXY* expression target the ribosome, not all of these are necessarily substrates of the pump (e.g., chloramphenicol). The mechanism underpinning this appears (at least, in part) to be integrally linked with ArmZ expression. The *armZ* ORF is preceded by a short leader peptide, whose translation leads to the formation of a specific mRNA secondary structure. In this structure, a transcriptional terminator is exposed prior to the RNA polymerase reaching the *armZ* ORF, thereby preventing *armZ* (and thence, *mexXY*) expression. However, when translation of the leader peptide is impaired, an alternative mRNA secondary structure forms in which the termination signal is occluded, allowing *armZ* expression (Morita *et al.*, 2009). It requires no great leap of the imagination to infer that the impaired translation accompanying the *fusA1*^*P443L*^ mutation could lead to a similar read-through of the terminator signal, enabling increased *armZ* expression. It is possible that similar mechanism(s) are responsible for some of the other pleiotropic (but nevertheless, discrete) effects associated with the *fusA1*^*P443L*^ mutation.

Residue P443 (*P. aeruginosa* numbering) is conserved in >70% of FusA1 sequences across multiple phyla (Margus *et al*., 2011). The P443→L substitution in the FusA1 of EMC1 introduces a hydrophobic residue (leucine) in place of a “structure breaking” one (proline). This was anticipated to have potential conformational consequences affecting the function or stability of the protein. Consistent with this, we found that purified FusA1^P443L^ mutant protein was somewhat less thermostable than the wild-type protein (ΔT_m_ = 1.4ºC) and displayed an altered intrinsic fluorescence profile. EF-G undergoes a series of large conformational reconfigurations during its catalytic cycle, especially upon ribosomal engagement. The most dynamic movement occurs in domain IV (responsible for binding mRNA in the A-site of the 30S ribosomal subunit), and this dictates the rotation of domains III and V (Li *et al.*, 2011). The P443→L substitution is positioned in a loop region between an α-helix and a β-strand in domain III. This is likely to be functionally significant because this loop lies directly opposite the critical “switch-1” and “switch-2” motifs in domain I, which are essential for GTPase activity. During the GTPase reaction, the switch motifs undergo large conformational changes, bringing domain I within bonding distance of domain III (Li *et al.*, 2011; Vetter, 2014; Macé *et al.*, 2018). Indeed, Mace *et al*. have identified a specific interaction between the switch-2 GTPase motif and residue Arg465 in domain III of the EF-G from *Thermus thermophiles* (Macé *et al.*, 2018). Given the very close three-dimensional proximity of P443 to this critical residue (which corresponds to H465 in *P. aeruginosa* FusA1), it is likely that mutation of P443 interferes with the local protein conformation and impairs guanine nucleotide exchange. This is significant because ribosome translocation proceeds at a far slower rate in the absence of EF-G GTPase activity (Holtkamp *et al.*, 2014) and although less is known about the role(s) of the EF-G GTPase in ribosome recycling (the likely main function of FusA1), the elevated levels of SmpB in EMC1 do indicate increased ribosomal stalling.

Perhaps the most significant finding of the current work is that the *fusA1*^*P443L*^ mutant displayed elevated expression of the T3SS (at the transcriptomic and proteomic level) and elevated expression of the T6SS machinery (at the transcriptomic level). This was expected because T6S and T3S are thought to be reciprocally-regulated. This reciprocal regulation is linked with signalling through the Gac pathway, which controls levels of free RsmA (Hood *et al.*, 2010). Excess free RsmA promotes T3SS expression and depresses T6SS/exopolysaccharide biosynthetic gene expression. The lower expression levels of *rsmA* in EMC1 are fully consistent with the observed up-regulation of T6S gene expression, but are inconsistent with the up-regulation of all five T3SS operons. However, we note that in some circumstances, the two secretion systems may be co-regulated. For example, genes encoding the T3 and T6 systems have been reported to be up-regulated in a *sigX* mutant (Gicquel *et al*., 2013). In this regard, we note that *sigX* was slightly (1.3-fold) down-regulated in EMC1.

In summary, we have shown here that a mutation in the gene encoding a ribosome recycling factor, FusA1, can lead to large-scale, but discrete alterations in the physiology of *P. aeruginosa. FusA1* mutants have been recently documented to confer resistance to aminoglycoside antibiotics (a phenotype that we confirm here), although their wider impact on virulence has not been reported. Our data show that one such *fusA1* mutant displays greatly up-regulated expression of the T3SS machinery, as well as increased resistance to gentamicin. Taken together, our data indicate that these changes may be linked with sensing of diminished translational capacity in the *fusA1* mutant, although additional work is required to confirm this and establish a mechanism. Consistent with this notion, it is worth recalling that treatment of *P. aeruginosa* with sub-MIC azithromycin (a macrolide targeting ribosome function) has also been shown to increase expression of the T3SS genes (Gillis *et al*., 2005).

## Acknowledgements

This work was supported by studentship MR/K50127X/1 (to EAM) from the MRC DTP programme, and by two flexible supplement awards (also from the MRC) for training in proteomic and transcriptomic analysis. RCT was supported by a PhD studentship from Hughes Hall Cambridge. Elements of the biochemical and genetic characterisation were supported by a grant from the Evelyn Trust.

## Materials and Methods

### Growth conditions

Unless otherwise stated, *P. aeruginosa* PAO1 strains were grown at 37°C in M9 minimal media supplemented with 0.5% (w/v) glucose. Planktonic cultures were grown with vigorous aeration for the indicated time or to the indicated optical density. Late exponential cells were harvested at 7 hours.

### EMC0 and plasmid construction

EMC0 (the P*cdrAB*::*lacZ* reporter strain) was constructed by sub-cloning the PCR-amplified 320 bp upstream region of *cdrAB* in to mini-CTX-*lacZ*, immediately in front of the promoterless *lacZ* ORF. The reporter plasmid was introduced into PAO1 by electroporation and transformants were selected for on 50 μg/mL tetracycline. Following integration of the construct into the chromosome, the mini-CTX backbone was removed by introducing pFLP2 into the transformants through bi-parental conjugation from β2163 (pFLP2). Transconjugants were selected for on 250 μg/mL carbenicillin. Transformants were streaked onto LB agar supplemented with 5% (w/v) sucrose to select for derivatives that had lost pFLP2, and then onto LB agar supplemented with 50 μg/mL tetracycline to verify successful excision of the tetracycline-resistance cassette. Loss of pFLP2 was confirmed through carbenicillin sensitivity. The resulting EMC0 was confirmed by PCR (and also by the whole genome sequence data).

The complementation vectors, p*fusA1* and p*fusA1*^*P443L*^, were constructed by PCR-amplifying the wild-type *fusA1* and mutated *fusA1*^*P443L*^ ORFs from PAO1- and EMC1-derived genomic DNA, respectively. The amplicons were then cloned into the PstI/HindIII sites in the MCS of pUCP20 (downstream of the *lac* promoter on the plasmid). The resulting plasmids were introduced into the recipient strains by electroporation and selection on 250 μg/mL carbenicillin. All plasmid constructs were confirmed by sequencing.

### Plasposon mutagenesis

The pTn*Mod*-OGm plasposon (Dennis *et al.*, 1998) was introduced into EMC0 *via* tri-parental mating. EMC0 was spotted onto solid agar with a helper *E. coli* strain (HB101 (pRK21013)) and *E. coli* JM109 (pTn*Mod*-OGm) for 18 hours. The mixed colony was then resuspended and transposon mutants were selected on agar media supplemented with 50 μg/mL^-1^ gentamicin and 30 μg/mL^-1^ X-Gal. After 30 hours, gentamicin resistant colonies were transferred to fresh gentamicin-supplemented plates and grown for a further 24 hours. EMC1 was isolated as a pale blue colony at this stage.

### β-galactosidase activity

Aliquots (100 μL) of planktonic culture were harvested over a 10 hour growth period and frozen at −80°C in a 96-well microtitre plate. After collection was complete, the plate was defrosted for 30 min at 37 °C and 10 μL were transferred to a fresh 96-well microtitre plate. The samples were frozen at −80°C for a second time, followed by thawing at room temperature. To quantitatively measure the level of β-galactosidase production, 100 μL of PBS containing 20 mg/mL lysozyme and 250 μg/mL 4-methylumbelliferyl-β-galactoside was added to the cells. The reaction progress was monitored every 30 sec for 30 min at 37°C in a Gemini XPS fluorimeter (Molecular Devices) using an excitation wavelength of 360 nm and emission wavelength of 450 nm.

### Whole genome sequencing

EMC0 and EMC1 were sequenced using Illumina technology (HiSeq 2500 platform) by MicrobesNG, Birmingham. Genomic DNA libraries were prepared using Nextera XT Library Prep Kit (Illumina) following the manufacturer’s protocol with the following modifications: two nanograms of DNA were used instead of one, and PCR elongation time was increased to 1 min from 30 seconds. DNA quantification and library preparation were carried out on a Hamilton Microlab STAR automated liquid handling system. Pooled libraries were quantified using the Kapa Biosystems Library Quantification Kit on a Roche light cycler 96 qPCR machine. Libraries were sequenced on the Illumina HiSeq using a 250bp paired end protocol. Reads were adapter trimmed using Trimmomatic 0.30 with a sliding window quality cutoff of Q15. *De novo* assembly was performed on samples using SPAdes version 3.7, and contigs were annotated using Prokka 1.11. Genome alignment and analysis was carried out using Mauve Multiple Genome Alignment (Darling *et al.*, 2004).

### Motility assays

To detect twitching motility, 10 mL of 1.5% (w/v) agar containing LB or M9 minimal media/glucose was prepared in a 10 cm diameter petri-dish. Colonies were stabbed into the agar with a sterile toothpick and the twitch halo was visualised after 24 hours of incubation at 37°C. To assess swimming motility, 25 mL LB or M9 minimal media/glucose containing 0.3% (w/v) Bacto agar was dispensed into a 10 cm diameter Petri-dish and allowed to solidify/surface dry at room temperature for 30 min. Bacterial cultures were normalised to an OD_600_ of 1, and 3 μL of culture was dispensed as a stab column into the agar using a pipette. The halo of swimming bacteria was assessed after 8-18 hours incubation at 37°C.

### Exoenzyme secretion assays

Gelatinase activity was measured on solid media plates containing 13 g/L nutrient broth and 30 g/L gelatin. Bacterial cultures were normalised to an OD_600_ of 1, and 5 μL was spotted onto the agar and left to soak in. Plates were incubated at 37°C overnight. Gelatinase activity was visualised by flooding the plates with saturated ammonium sulfate solution for 15 min to reveal the proteolytic halo.

### Exopolysaccharide secretion

Agar plates for the detection of exopolysaccharide production were prepared using 37 g/L Brain Heart Infusion broth supplemented with 50 g/L sucrose, and 0.8 g/L Congo Red. Bacterial cultures were normalised to an OD_600_ of 1 and 10 μL was spotted onto the surface of the agar. The plates were incubated at 37°C for 24 h. Exopolysaccharide production was determined semi-qualitatively through the amount of red pigmentation associated with the colony.

To quantitatively assess exopolysaccharide production, bacterial cultures were sub-cultured into fresh growth media supplemented with 10 μg/mL Congo Red and incubated at 37°C for 24 h on a rotating drum. The OD_600_ was measured and the cells were pelleted at 3200 x *g* for 10 min, at room temperature. This depletes any Congo Red bound to cell-associated polysaccharides; the greater the amount of polysaccharide sedimented, the lower the amount of Congo Red remaining in the culture supernatant. The optical density of the supernatant was measured at 495nm. The A_495_ was then normalised against the original OD_600_ of the culture.

### Siderophore detection

Bacterial cultures were harvested at late exponential phase. The cells were sedimented (3200 x *g* for 10 min) and the presence of siderophores in the cell-free culture supernatant was measured using the colorimetric SideroTech kit (Emergen Bio Inc.). A colour change occurs as ferric iron in the kit reagent binds to siderophores present in the culture supernatants.

### Western blotting

Following SDS-PAGE, proteins were transferred onto a polyvinylidene difluoride (PVDF) membrane using a Trans-Blot Turbo Transfer System (BioRad) with Trans-Blot Mini Transfer Packs. The membrane was incubated in blocking buffer (PBS containing 0.1% (v/v) Tween 20 and 5% (w/v) semi-skimmed milk powder) overnight, and then washed once in wash buffer (PBS and 0.1% (v/v) Tween 20). The membrane was then incubated for 1 hr with a primary anti-PcrV antibody (1:5000) in wash buffer. The membrane was then washed four times in wash buffer (5 min each wash), before the addition of IRDye 680RD (Li-Cor) secondary antibody. This was incubated for 45 min. The membrane was washed four times, as before and the protein bands were detected using an Odyssey CLx imaging system (Li-Cor).

### Proteomic analyses

Strains were grown with good aeration at 37°C to late exponential phase (OD_600_ of 0.6 – 0.8) in M9 minimal media supplemented with 0.5% (w/v) glucose. To ensure retention of the complementation and empty-vector plasmids, carbenicillin (250 μg/mL) was included in the starter culture, but not in the subculture used for proteomic analysis. We independently confirmed that the pUCP20 plasmid was retained by essentially all cells in the absence of selection over this sampling period (*data not shown*). Cells from 45 mL culture were harvested at 3200 x *g* for 30 min at 4°C. The cell pellet was resuspended in PBS and sedimented a second time. Pellets were then resuspended in 800 μL of lysis buffer (100 mM Tris-HCl, 50 mM NaCl, 20 mM EDTA, 10% (v/v) glycerol, 1 mM DTT, pH 7.5) containing a Complete™ protease inhibitor cocktail tablet (Roche), and sonicated (3 x 5 sec at 15 amps, MSE microtip) on ice. Unlysed cells and debris were pelleted at 21,000 x *g* for 30 min at 4°C. The protein concentration of the supernatant was determined using the DC protein assay (Biorad). LC-MS/MS was performed by the Cambridge Centre for Proteomics. The samples were digested with trypsin and dried. The dried peptides were reconstituted in 100 mM triethylammonium bicarbonate and labelled using 10-plex TMT (tandem mass tag) reagents according to the manufacturer’s (Thermo Scientific) protocol. Tagged peptides were fractionated by reverse-phase chromatography and were identified and quantified using a high resolution Orbitrap mass spectrometer coupled to a Dionex Ultimate 3000 RSLC nano UPLC (Thermo Fischer Scientific). Proteomic data sets were analysed with the empirical Bayes moderated T-test implemented by the limma package (Ritchie *et al.*, 2015). P-values were corrected for multiple hypothesis testing using the Benjamini-Hochberg method (FDR ≤ 0.05). Differential expression was calculated based on normalized log_2_ ratios. The MS/MS fragmentation data was searched against the National Centre for Biotechnology Information (NCBI) database using MASCOT (Matrix Science) search engine.

### Expression and purification of FusA1

For FusA1 purification, the wild-type and mutated *fusA1* genes were PCR-amplified (from PAO1 and EMC1, respectively) and cloned into pET-19m. The resulting plasmids were introduced into Rosetta *E. coli* cells. The Rosetta cultures were grown up in 1 L of LB supplemented with 50 μg/mL carbenicillin, and incubated at 37°C to an OD_600_ of 0.6 - 0.7. Isopropyl-β-D-thiogalactopyranoside (IPTG) was added to the cultures to a final concentration of 1 mM and the culture was incubated at 20°C for 24 hours. Cells were centrifuged at 3430 x *g*, for 20 min at 4°C. The cell pellet was resuspended in 10 mL lysis buffer ((100 mM Tris-HCl, 50 mM NaCl, 20 mM EDTA, 10% (v/v) glycerol, 1 mM DTT, pH 7.5) containing a Complete™ protease inhibitor cocktail tablet (Roche)) and lysed by sonication (5 x 30 sec at 13 amps) on ice. The samples were then clarified by centrifugation (14,636 x *g* for 30 min, 4°C), and the supernatant was passed through a 0.45 μm membrane filter (Millipore). FusA1 protein was purified from the soluble, filtered extracts by loading onto a pre-equilibrated Ni-NTA Superflow Cartridge (Qiagen) at 4°C. The column was equilibrated with protein purification buffer (50 mM sodium phosphate, 200 mM NaCl, 10 mM imidazole, 10% (v/v) glycerol, pH 8.0). The column was washed until the A_280_ of the eluate was negligible, and eluted in the same buffer containing 250 mM imidazole. The eluted protein was concentrated in a Vivaspin-20 and then dialysed against 1 L of 50 mM Tris, 100 mM NaCl, 5% (v/v) glycerol, pH 7.4). Aliquots of the protein were stored frozen at −80°C until use.

### Intrinsic tryptophan fluorescence

The concentration of appropriately diluted purified His_6_FusA1 was determined spectrophotometrically at 280 nm using a quartz cuvette (1cm pathlength) and assuming MW = 78738 and ε = 61310 M^-1^cm^-1^. EF-G protein was diluted in 2 mL dialysis buffer to a final concentration of 0.8 μM. Intrinsic Trp fluorescence was measured in a thermostated (25°C) quartz cuvette, on a FP-8300 Spectrofluorometer (JASCO) using an excitation wavelength of 295 nm and an emission wavelength of 305 – 400 nm (0.5 nm intervals, 100 nm/min). All spectra were recorded in triplicate and then averaged.

### Thermal shift analyses

Purified wild-type FusA1 or FusA1^P443L^ (20 μL volume containing 20 μM protein) were mixed with SYPRO Orange dye and subjected to a thermal scan (25ºC → 95ºC) in a Roche Lightcycler 480, and the fluorescence was measured at 483 nm and 568 nm.

### RNA extraction and sequencing

EMC0, EMC1 (both containing the pUCP20 empty vector) and EMC1* (containing pUCP20:*fusA1)* were cultured in M9 minimal media supplemented with glucose and samples were harvested at the late exponential phase of growth (OD_600_ of 0.6 – 0.8) into RNA Later (Ambion). The samples were incubated at 4°C for 15 min and pelleted at 21,000 x *g* for 20 min at 4°C. Total RNA was extracted using the RNeasy Mini Kit (Qiagen) and digested twice with on-the-column DNase I digestion, following the manufacturer’s guidelines. The concentration and purity of the resulting RNA was measured using a NanoDrop ND-1000 Spectrophotometer. The absence of contaminating proteins and organic compounds was indicated by an A_260/280_ ratio of 1.8 – 2.0 and A_260/230_ nm of 2.0 – 2.2. Total RNA was sent to GATC Biotech (De) for rRNA depletion and RNA-Sequencing using the Illumina platform (10 million reads per sample, single read, 1 x 50 bp). The RNA-Seq reads were processed using FastaQC and were mapped to the PAO1 genome and analysed using the Tuxedo Suite package. The principal component analysis plot and the volcano plots were constructed in R.

For the Q-RT-PCR analyses, complementary DNA (cDNA) synthesis was required. To do this, 1 μg of RNA was combined with 50 ng oligo(dT)_15_, 300 ng random hexamers, and 1 μL dNTP mix. The samples were then heated at 65°C for 5 min followed by incubation on ice for 2 min. To this mixture, 4 μL of 5 x SuperScript III First strand buffer, 1 μL 0.1 M DTT and 1 μL SuperScript III Reverse transcriptase (Invitrogen) was added. Samples were incubated at 25°C for 5 min, followed by 50°C for 60 min. The reaction was inactivated by heating to 70°C for 15 min. cDNA was used as a template for RT-PCR using 20 - 25 cycles of 98°C (10 sec), 55°C (20 sec), 72°C (20 sec/kb).

