## Supplementary material for "Global reprogramming of virulence and antibiotic resistance in *Pseudomonas aeruginosa* by a single nucleotide polymorphism in the elongation factor-encoding gene, *fusA1*": SI data

**Supplementary Figures and Tables**

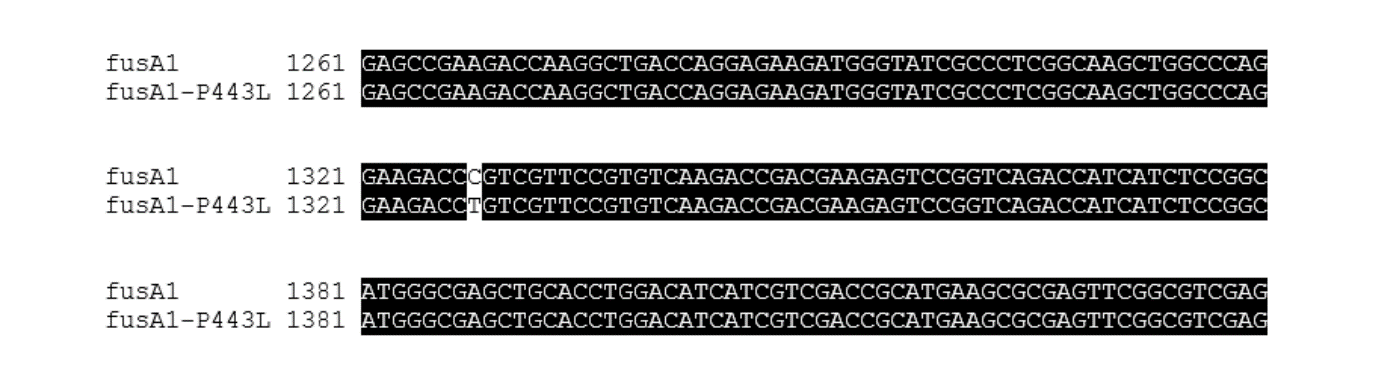

**Figure S1**. **Alignment of wild-type *fusA1* from EMC0 with *fusA1* from EMC1**. The alignment shows the cytosine to thymidine transition that leads to the P443L substitution in the amino acid sequence.

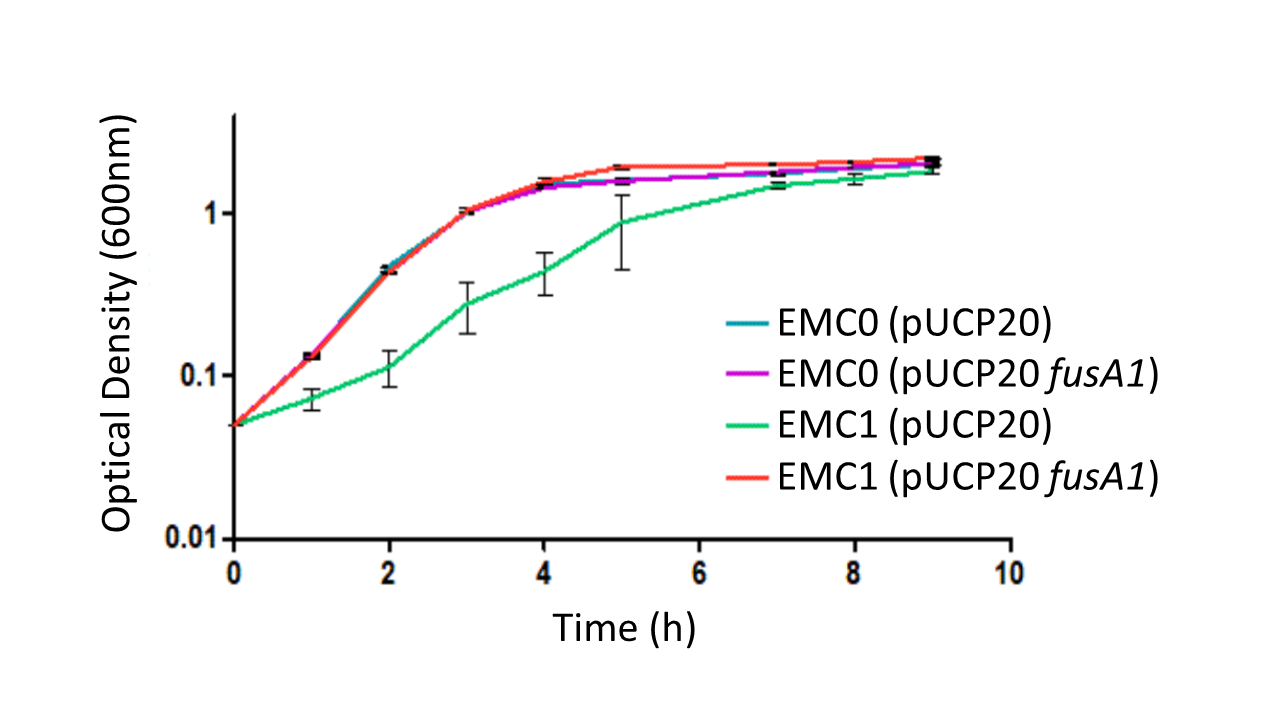

**Figure S2**. **The ECM1 growth defect in LB is complemented by expression of wild-type *fusA1 in trans***. The figure shows growth curves (monitored as OD_600_) of EMC0 (blue, purple) and EMC1 (green, red) containing pUCP20 (empty vector control plasmid) or pUCP20 with cloned wild-type *fusA1*. Transcription of the cloned wild-type gene was driven by the *lac* promoter on pUCP20. The cloned ORF was engineered to contain an optimized ribosome binding site (AGGAGGT) 5 nucleotides upstream of the GTG start codon, thereby ensuring high-level expression.

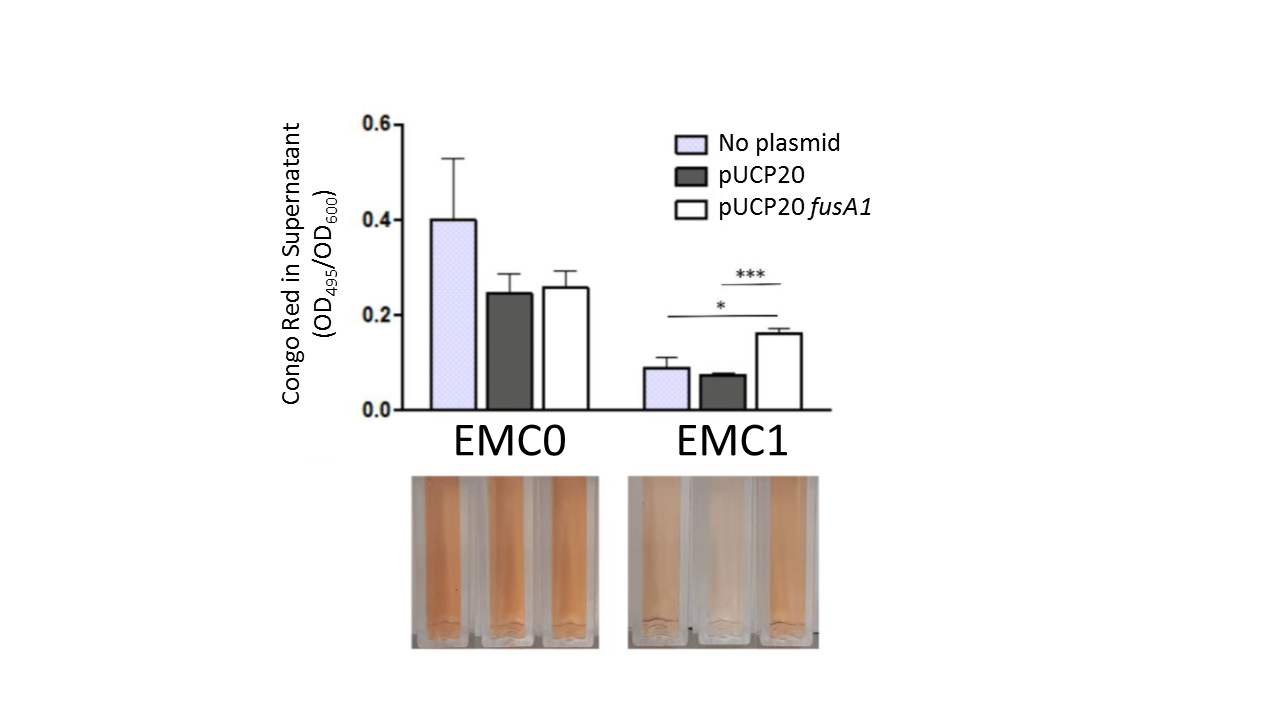

**Figure S3. Quantitative Congo Red assay for the production of exopolysaccharides in planktonic culture**. To quantitatively assess exopolysaccharide production, overnight bacterial starter cultures were inoculated in fresh growth media supplemented with 10 μg/mL Congo Red. Cultures were grown with good aeration at 37ºC for 24 h on a rotating drum. The OD600 was measured and the cells were pelleted at 3200 x *g* for 10 min at room temperature. The principle behind this is that any Congo Red bound to cell-associated polysaccharides becomes sedimented out of the solution. Unbound Congo Red in solution was determined by quantifying the optical density of the supernatant at 495 nm. The OD495 was normalised against the culture OD600. The greater the amount of polysaccharide produced, the lower the final A_495_ of the culture supernatant. Statistical significance between groups was assessed by an unpaired *t*-test (n=3) (* = *p* <0.05, ** = *p* <0.01).

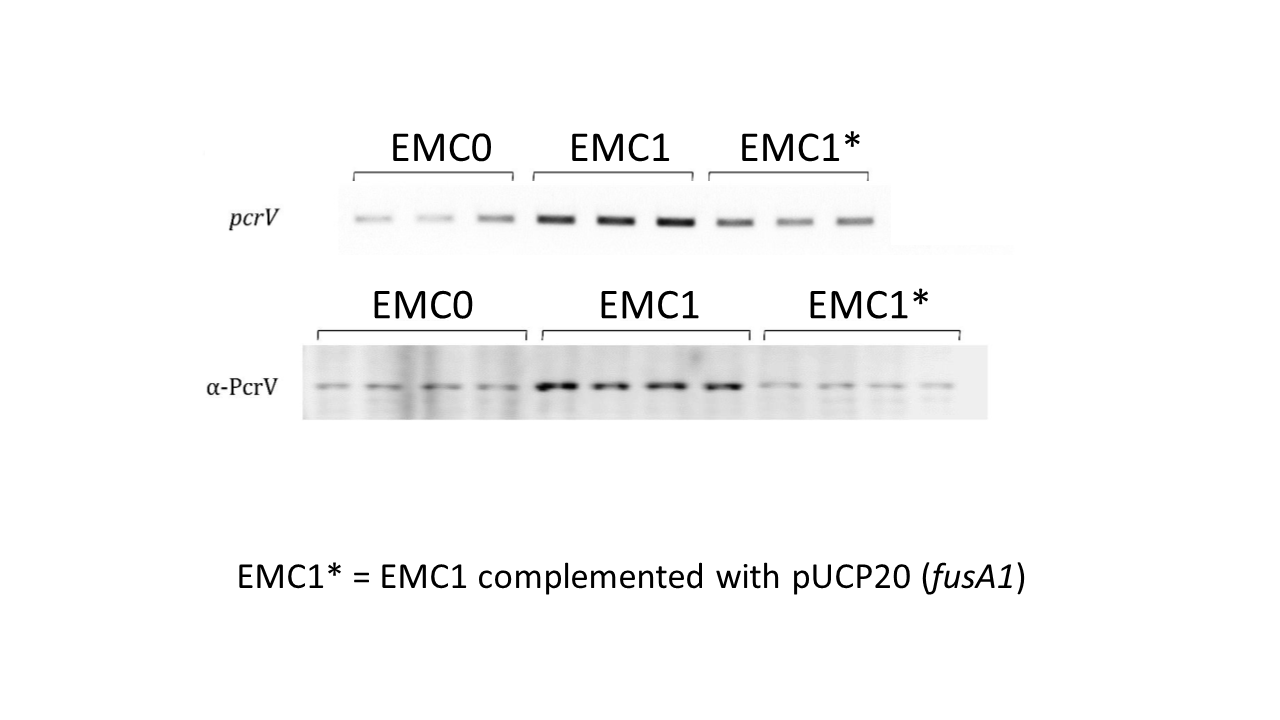

**Figure S4**. **Expression of the Type III Secretion machinery protein, PcrV, is stimulated in EMC1**. Upper panel; RT-PCR amplification the *pcrV* ORF (25 cycles of PCR amplification). Lower panel; western blot using α-PcrV antibodies on whole cell protein samples of each strain. Samples were taken for analysis from independent triplicate (RT-PCR) or quadruplicate (western) cultures. The EMC0 and EMC1 samples contained empty pUCP20 vector.

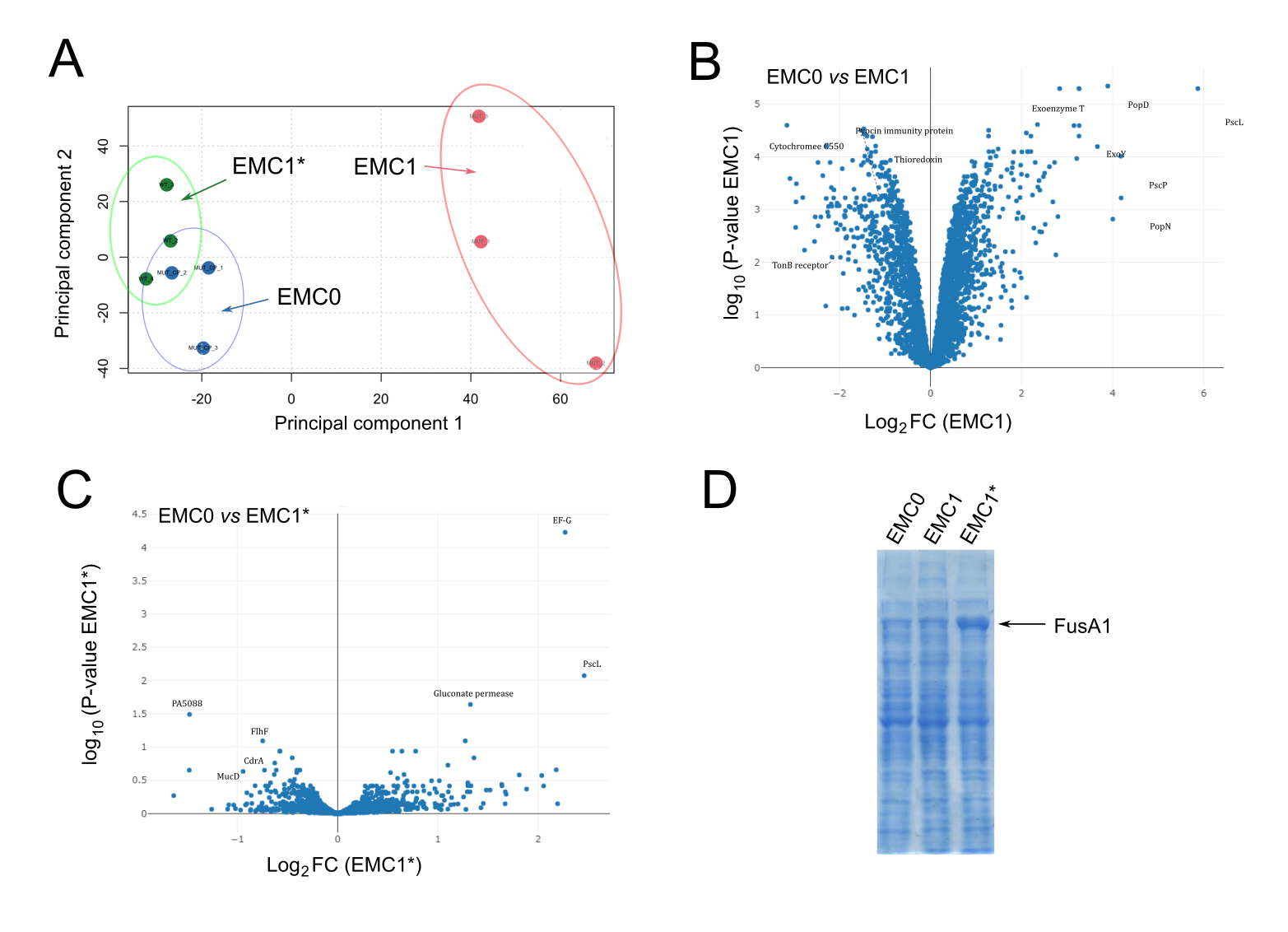

**Figure S5. The *fusA1^P443L^* mutation leads to global proteomic changes in EMC1**. **(A)** Principal Components Analysis (PCA) scores plot showing the global trends observed in EMC0 blue circles), EMC1 (red circles) and EMC1* (green circles). **(B)** “Volcano plot” showing log_2_FC of each identified protein *vs* P-value for the comparison of proteins in EMC1 and EMC0. (C) Volcano plot showing log_2_FC of each identified protein *vs* P-value for the comparison of proteins in EMC1* and EMC0. In (B) and (C) a selection of the identified proteins are indicated on each plot. Note that in (C), FusA1 (EF-G) is the most highly-modulated protein. This is because FusA1 has been over-expressed from the pUCP20 *fusA1* used to complement the *fusA1^P443L^* mutation in EMC1. The over-expressed FusA1 protein band can be clearly seen in the 1D SDS-PAGE profile of the EMC1* in panel **(D)**.

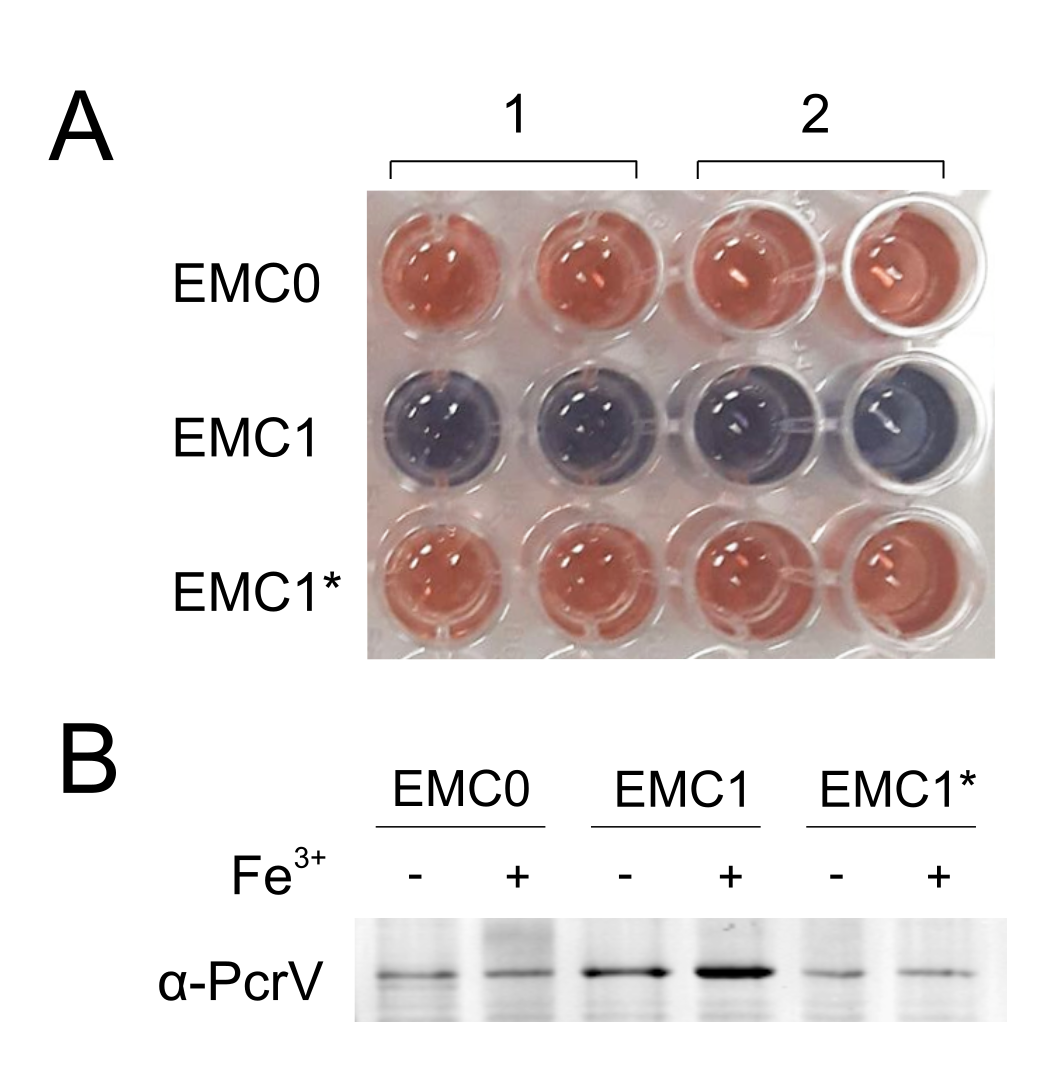

**Figure S6. Siderophore production by EMC0, EMC1 and EMC1***. **(A)** Siderotech assay was used for the detection of siderophores in the culture supernatants of EMC0, EMC1 and EMC1*. The supernatant from two biological replicates (1 and 2) were tested twice for each indicated strain. The solution turns from blue to red upon the detection of siderophores in the culture supernatant. **(B)** Western blot showing whole cell lysates probed with α-PcrV antibodies. The indicated strains were grown in M9 minimal media ± 100 μM FeCl_3_.

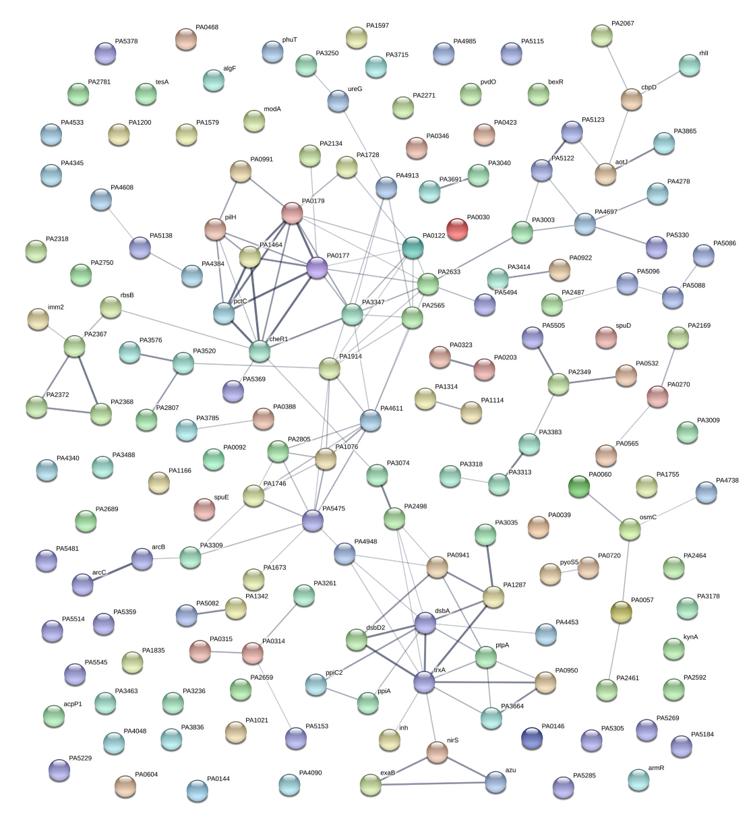

**Figure S7. STRING analysis of proteins that were down-regulated in EMC1 compared with EMC0**. A selection of the more highly-interconnected clusters of proteins are discussed in the main body text.

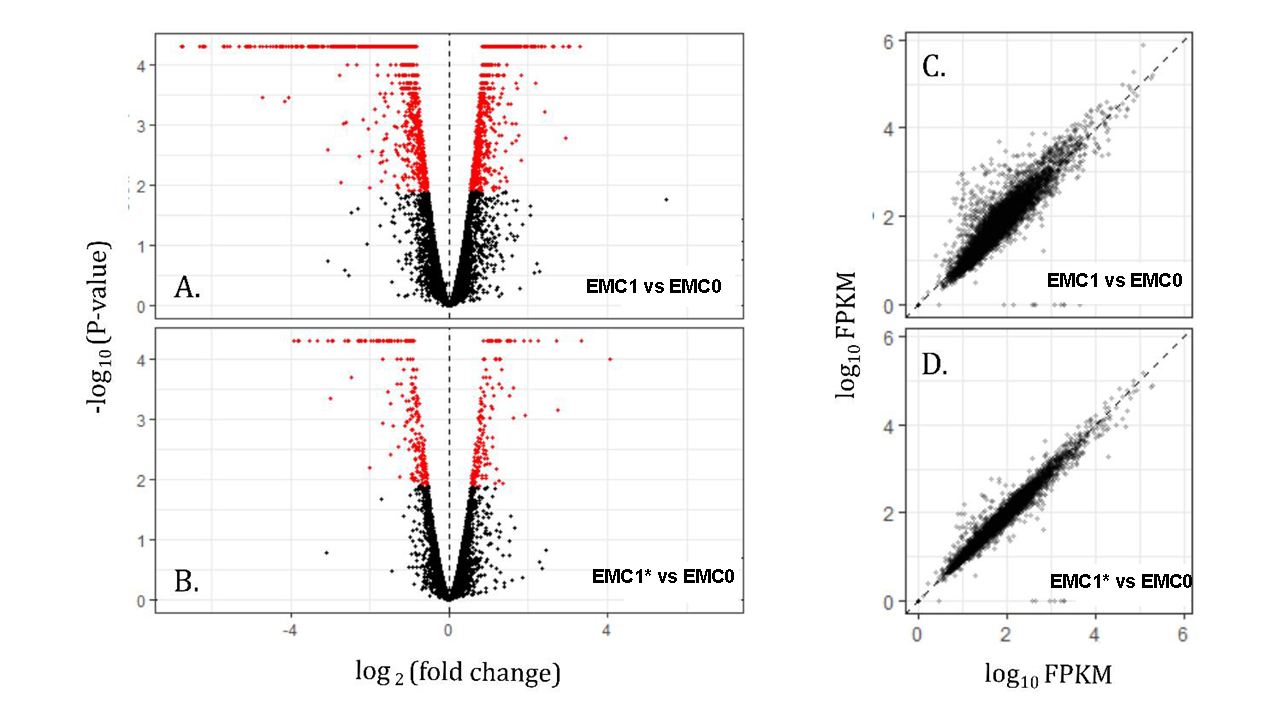

**Figure S8. Volcano plot representing the log2 fold change (FC) of RNA transcript abundance against statistical significance for (A) EMC1, and (B) the EMC1* in relation to the progenitor strain, EMC0.** Significantly (P < 0.01) modified transcripts are highlighted in red. Note that far fewer proteins are modulated in (B) compared with (A). An FPKM (fragments per kilobase, per million) scatter plot comparing the log10 ratios of FPKM expression values for the progenitor strain with **(C)** EMC1, and **(D)** EMC1*.

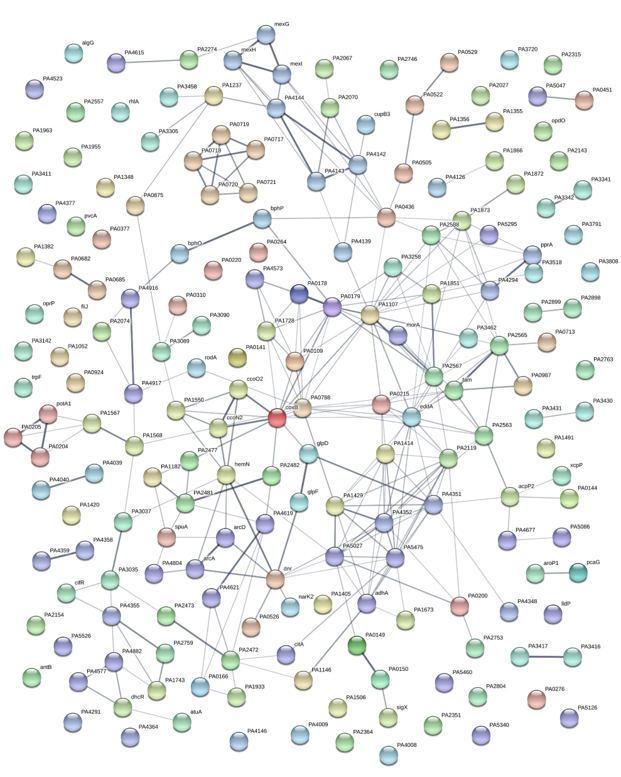

**Figure S9. STRING analysis of transcripts that were down-regulated in EMC1 compared with EMC0**. A selection of the more highly-interconnected clusters are discussed in the main body text.

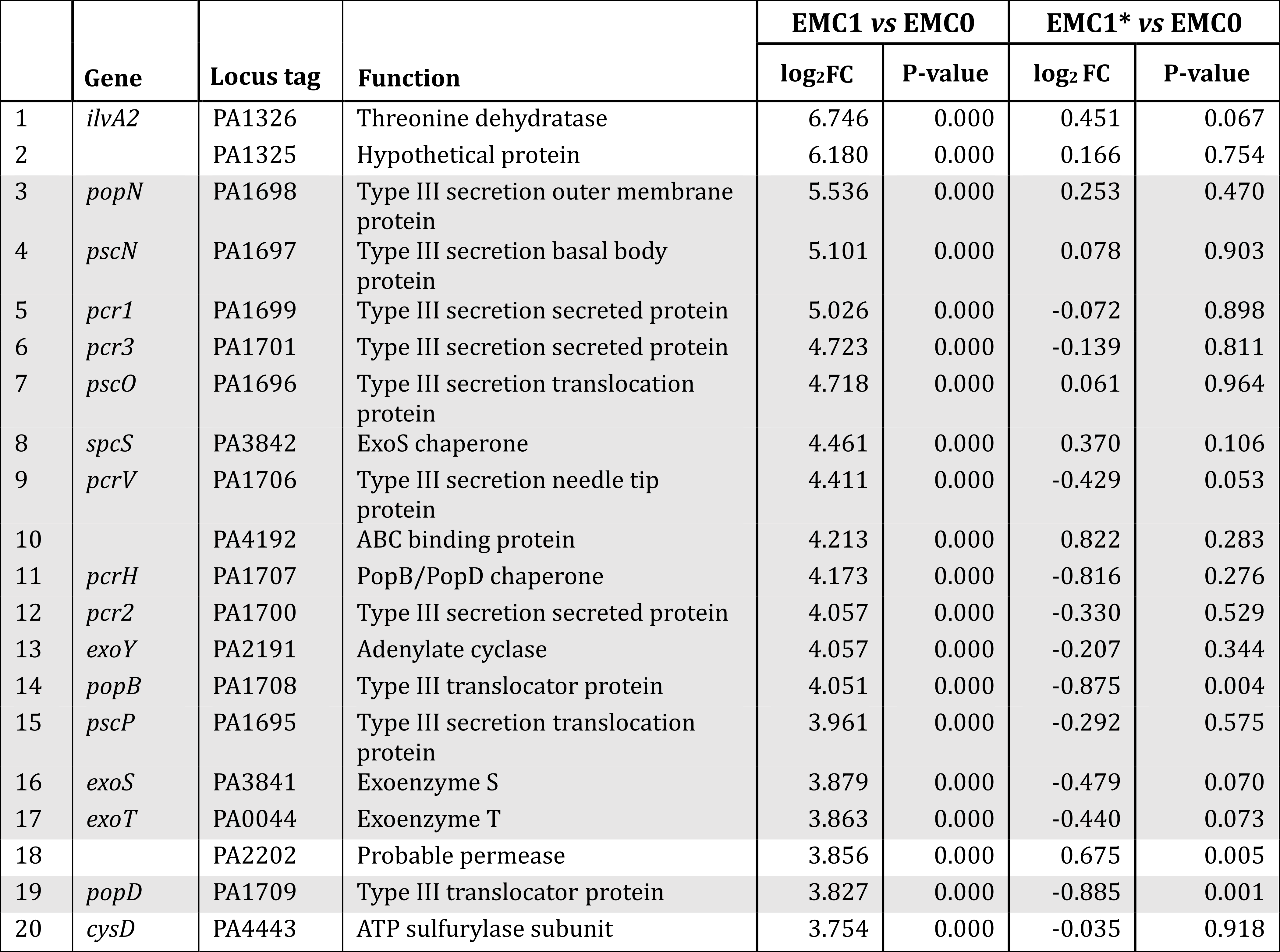

**Table S1. Expression of the 20 mostly highly up-regulated transcripts in EMC1 is reversed by expression of wild-type *fusA1 in trans* in EMC1***. Proteins involved in T3S are shaded.
